# Differential proteomic analysis of laser-microdissected penetration glands of avian schistosome cercariae with a focus on proteins involved in host invasion

**DOI:** 10.1101/2021.08.24.457505

**Authors:** Oldřich Vondráček, Libor Mikeš, Pavel Talacko, Roman Leontovyč, Jana Bulantová, Petr Horák

## Abstract

Schistosome invasive stages, cercariae, leave intermediate snail hosts, penetrate the skin of definitive hosts, and transform to schistosomula migrating to final localization. During invasion, cercariae employ histolytic and other bioactive products of specialized holocrine secretory cells – postacetabular (PA) and circumacetabular (CA) penetration glands. Although several studies attempted to characterize protein composition of the *in vitro* induced gland secretions in *Schistosoma mansoni* and *Schistosoma japonicum*, the results were inconsistent and dependent on the method of sample collection and processing. Products of both gland types mixed during their secretion did not allow localization of identified proteins to a particular gland. Here we compared proteomes of separately isolated cercarial gland cells of the avian schistosome *Trichobilharzia szidati* employing laser-assisted microdissection and shotgun LC-MS/MS, thus obtaining the largest dataset so far concerning the representation and localization of cercarial penetration gland proteins. We optimized the methods of sample processing with cercarial bodies (heads) first. Alizarin-pre-stained, chemically non-fixed samples provided optimal results of MS analyses, and enabled distinguishing PA and CA glands for microdissection. Using 7.5 × 10^6^ μm^3^ sample volume per gland replicate, we identified 3347 peptides assigned to 792 proteins, from which 461 occurred in at least 2 of 3 replicates in either gland type (PA = 455, 40 exclusives; CA = 421, 6 exclusives; 60 proteins differed significantly in their abundance between the glands). Peptidases of 5 catalytic types accounted for ca. 8 % and 6 % of reliably identified proteins in PA and CA glands, respectively. Invadolysin, nardilysin, cathepsins B2 and L3, and elastase 2b orthologs were the major gland endopeptidases. Two cystatins and a serpin were highly abundant peptidase inhibitors in the glands. CA glands were rich in venom allergen-like proteins. The assembled total cercarial body proteome included 1631 identified proteins and revealed additional interesting factors possibly related to tissue invasion.

**Highlights:**

- Proteomes of two penetration gland types in schistosome cercariae greatly differ
- Postacetabular glands possess 40 unique proteins and are abundant in hydrolases
- Circumacetabular glands posses 6 unique proteins and are rich in VAL proteins
- Peptidases make up 8 % of postacetabular and 6 % of circumacetabular gland proteins
- Cercarial elastase is unique to circumacetabular glands of *Trichobilharzia szidati*

Note: Supplementary data associated with this article All supplementary data files can be accessed from the following link: http://www.helminthology.cz/supplementary_files.html

## 1. Introduction

The determination of protein functions in an organism mostly requires accurate localization of the protein or, alternatively, of its coding mRNA. Classical and widely used approaches include laborious and often lengthy antibody-mediated immunolocalization or RNA *in situ* hybridization accompanied by possible inconveniences including, e.g., the necessity of using laboratory animals, low immunogenicity of some antigens, low specificity of antibodies, demanding preparation and processing of material for histology, exacting optimization of RNA probes and conditions of hybridization, and generally low throughput. In the post-genomic era, many of these obstacles can be solved by a combined approach employing laser microdissection (LMD) that allows to isolate the cells/tissues of interest from histological sections under constant microscopic supervision, and mass spectrometry (MS) that enables precise protein identification (e.g. Espina et al., 2007; Xu, 2010; Legres et al., 2014). Besides a broad spectrum of applications in biology and medicine, this approach has also been used in proteomic studies of parasites and host-parasite interactions in cases of, e.g., the causal agents of malaria (Swearingen et al., 2017; Lindner et al., 2019), the tissue-dwelling stages of the tapeworm *Echinococcus granulosus* (Longuespée et al., 2017) or the intramolluscan stages of the human blood fluke *Schistosoma mansoni* (Dinguirard et al., 2018).

In the present study, we applied LMD+MS to determine the protein contents of two types of peculiar glands in the invasive stage, cercaria, of an avian schistosome blood fluke *Trichobilharzia szidati* (Trematoda: Schistosomatidae). Schistosomes are important parasites of mammals or birds, depending on the parasite genus. The members of the genus *Schistosoma* cause a severe and potentially lethal disease (schistosomiasis, bilharziasis) affecting human health in 52 endemic countries, with at least 229 mil. people requiring preventive treatment in 2018 (WHO Schistosomiasis Fact Sheet, https://www.who.int/news-room/fact-sheets/detail/schistosomiasis). Cercariae of avian schistosomes can also penetrate the human skin and cause an allergic skin reaction termed cercarial dermatitis or swimmers’ itch (Horák et al., 2015; Macháček et al., 2018). Schistosomes have two-host life cycles including intermediate aquatic snail hosts and homoiotherm vertebrates as definitive hosts. The final stages of intramolluscan development, cercariae, escape into water. They actively penetrate the skin of their final hosts while releasing the contents of so called ‘penetration glands’ packed with histolytic peptidases and many other bioactive substances. In the same time they shed their tails and a thick glycocalyx, build a double tegumental membrane, and undergo dramatic physiological changes, thus transforming to juvenile worms (schistosomula) (Dvořák et al., 2008; Kašný et al., 2009; Horák et al., 2015; Gobert and Nawaratana, 2016; Kašný et al., 2016; Řimnáčová et al., 2017; Horák et al., 1998).

All known schistosome cercariae possess two types of these acetabular penetration glands in a typical arrangement. They are termed according to their position towards the ventral sucker (acetabulum) of the cercaria as postacetabular (PA) and circumacetabular (CA) (originally described as preacetabular) glands. Both types of the glands are unicellular, mesenchymal by origin, exocrine with holocrine type of secretion, but they differ in ultrastructure, morphology of secretory vesicles, composition, and affinity to various histological dyes (Stirewalt and Kruidenier, 1961; Bruckner, 1974; Fishelson et al., 1992; Mikeš et al., 2005; Ligasová et al., 2011). Three pairs of PA glands (PAg) are located largely behind the acetabulum and occupy a major part of cercarial hind body. Two pairs of CA glands (CAg) surround the base of acetabulum. In cercariae of the avian schistosome *Trichobilharzia regenti*, the penetration glands occupy more than 1/4 of cercarial body volume – PAg ca. 15 % and CAg ca. 12 % (Ligasová et al., 2011). Their secretions have been believed to be synthesized during intramolluscan development (Dorsey, 1975), since cercariae seem to be transcriptionally silent (Roquis et al., 2015), although they are equipped with (pre-synthesized?) mRNA (Leontovyč et al., 2016). The ducts of these unicellular glands are just cytoplasmic processes of the gland cells which lead to the apex of cercaria and release gland contents upon specific chemical stimuli from the host skin (Kašný et al., 2016).

Many studies aimed at identifying the content of schistosome penetration glands have been based on analyses of cercarial excretory/secretory products (ESP) after *in vitro* induction of gland emptying by mechanical or chemical stimuli, where the products of both gland types have been mixed and contaminated by somatic and tegumental proteins of cercariae, proteins of snail origin, and proteins from the experimental environment. It seems from the literature that the method (stimulus) used for induction of gland emptying greatly affects the results. For example, in *S. mansoni*, 53 parasite proteins were determined in human skin lipid-induced ESP separated by SDS-PAGE and analyzed by shotgun LC-MS/MS of the bands (Knudsen et al., 2005). Mechanical removal of the tail by vortexing *S. mansoni* cercariae and subsequent incubation of cercarial bodies in RPMI-1640 medium resulted in MS identification of 16 proteins from 50 most abundant protein spots after 2-D electrophoresis – 7 of vesicular origin and 9 cytosolic proteins (Curwen et al., 2006). In skin graft-induced ESP of *S. mansoni* analyzed by shotgun LC-MS/MS, around 100 proteins were identified, both vesicular and cytosolic (Hansell et al., 2008). In *Schistosoma japonicum* cercariae sheared through a 22 gauge needle to remove tails and start-up the transformation in SCN medium, the LC-MS/MS analysis identified 361 unique proteins in in-gel digested complete ESP after SDS-PAGE (Dvořák et al., 2008). A total number of 1972 proteins were identified in whole cercariae of *S. japonicum* using SDS-PAGE/LC-MS/MS analysis, including 46 cercarial proteases, from which 25 apparently participate in the penetration process (Liu et al., 2015).

Most of the published work focusing on analysis of cercarial penetration glands has been performed on human blood flukes. Little is known about the gland content in cercariae of avian schistosomes. In cercarial ESP of *Trichobilharzia szidati* and *T. regenti*, peptidase activities of predominantly cysteine catalytic type have been detected (Mikeš et al., 2005; Kašný et al., 2007). A cysteine peptidase cathepsin B2 was later localized in PAg of *T. regenti* and its ability to cleave proteins of the host skin was demonstrated (Dolečková et al., 2009). The presence of a lectin-like activity was reported in *T. szidati* PAg (Horák et al., 1997). Besides, a high content of Ca^2+^ in CAg, typical of schistosomes, was confirmed in both *T. regenti* and *T. szidati* (Modha et al., 1998; Mikeš et al., 2005).

Here we present for the first time differential proteomes of CAg and PAg from schistosome cercariae, opting for the avian schistosome *T. szidati* as a model species. The data were obtained by MS analysis of laser-dissected penetration gland cells. The methods have been experimentally optimized by evaluation of several factors having possible effect on the results, namely volume of the microdissected tissue, mode of tissue fixation, application of histological dyes, processing time vs. material degradation, and mode of processing of the microdissected material for MS analysis. We took the advantage of the availability of annotated *T. szidati* cercarial transcriptome and *T. regenti, S. mansoni*, and *S. japonicum* genomes for proteomic data analysis. Moreover, as a by-product of the MS analyses related to method optimizations, the total proteome of *T. szidati* cercarial body excluding tail is also presented.

## 2. Materials and Methods

### 2.1. Maintenance of parasite life cycle and ethics statement

The life cycle of the avian schistosome *T. szidati* has been routinely maintained in the Laboratory of Helminthology, Faculty of Science, Charles University using the intermediate snail hosts *Lymnaea stagnalis* and the domestic ducks *Anas platyrhynchos* f. dom. as the definitive hosts (Meuleman et al., 1984). The animals were treated in concordance with the legislation of the Czech Republic (246/1992 and 359/2012), the European Directive (2010/63/EU), and ARRIVE guidelines. The experiments were approved by the Professional Ethics Committee of the Faculty of Science, Charles University, and of the Research and Development Section of the Ministry of Education, Youth, and Sports of the Czech Republic (approval nos. MSMT-31114/2013-9 and MSMT-33740/2017-2). The animal facility, its equipment, animal welfare, and accompanying services, including the maintenance of experimental animals, have been approved by the Section of Animal Commodities of the Ministry of Agriculture of the Czech Republic (approval no. 13060/2014-MZE-17214).

### 2.2. Collection of cercariae

Cercariae were released from infected snails within an hour after illumination in glass beakers with dechlorinated tap water. The suspension of cercariae was transferred to long-necked flasks wrapped in aluminium foil and illuminated from above. Parasites concentrated under the water surface due to their positive phototaxis were collected, cooled in a 50 ml Falcon tube on ice till they dropped to the bottom, transferred to micro tubes, centrifuged (2000 × g, 2.5 min, 1 °C), and separated from the excess water to obtain a dense suspension of living cercariae.

### 2.3. Embedding and cryosectioning

Cercariae were transferred into polyethylene embedding capsules (Conical Tip Capsules, BEEM) and centrifuged in a cooled centrifuge (2000 × g, 2.5 min, 1 °C). Finally, cercarial pellets were mixed with Tissue Freezing Medium (Leica) and frozen at –80 °C until use. Sections of 10 μm were prepared in a cryostat (CM3050 S Research Cryostat, Leica) and mounted onto membrane frame slides (MMI). The slides were stored at –80 °C until use. In the case of histological staining, the stained cryosections were washed with dH_2_0. All cryosections were air-dried for 10 min at room temperature (RT) prior to laser microdissection.

### 2.4. Laser microdissection

Microdissections of areas of interest were carried out using the MMI SmartCut system (Olympus CKX41 inverted microscope; 40x microdissection objective lens; Olympus SmartCut Plus software). The microdissects were recovered mechanically by transparent silicone 0.5 ml Isolation Caps (MMI). Tubes with collected dissects were stored at –80 °C until MS analysis.

### 2.5. Optimization of the amount of material and the method of sample preparation for MS analysis

We performed MS analyses of various volumes of microdissected cercarial tissues in order to find an optimal value (limiting volume) for a solid MS analysis, trying to get a high number of detected proteins on the one hand, and minimize the labour intensity and demands on time on the other hand. Cryosections from non-fixed cercariae were used. The areas of entire cercarial bodies (including penetration glands but without tails) were microdissected. Volumes of 2.5, 5, 7.5, 10, and 12.5 × 10^6^ μm^3^ were tested. Two biological replicates were collected for each volume. Three different methods of sample preparation for MS analysis were tested with all the above-mentioned sample volumes: In-solution digestion (ISD), In-StageTips (IST) and “SP3” (SP3) (protocols to be seen below). Based on the best results obtained with the IST method, this protocol was re-tested on an extended range of sample volumes using 1, 2.5, 5, 7.5, 10, and 30 × 10^6^ μm^3^ of microdissected tissues. Two biological replicates were taken for each volume in this case.

### 2.6. Optimization of tissue preservation protocol

In order to find an optimal protocol for processing of the cercariae, three approaches have been applied: living cercariae were a) used in a native state without fixation (NF samples), b) fixed with 70% ethanol for 20 minutes, then thoroughly washed 5 × 5 min in distilled water (dH_2_O) to remove ethanol (EtOH samples), and c) fixed with freshly depolymerized 4% paraformaldehyde for 1 h, then washed 3 × 5 min in PBS (PF samples).

### 2.7. Optimization of staining protocol for penetration glands and cercarial tissues

In order to optimize the protocol for cell staining on cryosections and to evaluate the impact of several histological dyes on proteomic analysis, the cercariae were stained by selected dyes in two ways: 1) prior to freezing of living cercariae or 2) on thawed cryosections of parasites, just prior to laser microdissection.

#### 2.7.1. Staining of cercariae before cryopreservation

Samples of concentrated living cercariae were stained with saturated aqueous solution of alizarin (Merck) aded to a suspension of cercariae at the v/v ratio of 2:1 for 15 min (last 5 minutes on ice), and washed quickly with water (4 °C).

#### 2.7.2. Staining of cryosections of cercariae

Cryosections of non-fixed cercariae were thawed for 10 min. The cryosections were further processed in four ways: a) left unstained, b) stained for 2 min with 1% toluidine blue, c) stained for 10 min with saturated lithium carmine solution diluted 1:4 with 70% ethanol, or d) stained for 10 min with 0.2% trypan blue diluted 1:4 with 70% ethanol.

### 2.8. Microdissections of cercariae used for evaluation of the fixation/staining procedures

The areas of entire cercarial bodies (including penetration glands but without tails) were microdissected from the cryosections. Six biological replicates for every type of tissue preservation method (NF, EtOH, and PFA, each sample containing 7.5 × 10^6^ μm^3^ of cercarial tissue) were prepared for MS analysis (further processed as 6 technical replicates for each fixation method). All samples were frozen at –80 °C. Five biological replicates were prepared for every type of staining (non-stained, alizarin, toluidine blue, lithium carmine, trypan blue), each containing 7.5 × 10^6^ μm^3^ of cercarial tissues (further processed as 5 technical replicates for each staining method).

### 2.9. The effect of processing time on sample stability

In an attempt to test possible effect of prolonged microdissection at room temperature (RT) on subsequent proteomic analysis, cryosections of non-fixed cercariae were used. A total of 12 samples were microdissected (areas of whole cercarial bodies excluding tails), each of 7.5 × 10^6^ μm^3^. All samples were then exposed to room temperature for 1, 2, 3, 5, 8, and 24 hours including the microdissection process (two biological replicates for each time interval). Thereafter, the samples were boiled in a lysis buffer (see IST protocol below) and then frozen at –80 °C. Thawed samples were further processed according to the IST method.

### 2.10. Microdissections of cercarial penetration glands

Microdissections of areas from CAg and PAg were carried out from cryosections of cercariae stained with alizarin prior to cryopreservation (see above). In schistosome cercariae, alizarin stains intravitally and selectively the content of CAg due to a high content of Ca^2+^ (Mikeš et al., 2005; Ligasová et al., 2011). Thus, gland cells labeled with alizarin were isolated as CAg. PAg distinguishable from the other tissue thanks to the light refraction were isolated only when all of the following conditions were met: 1) Dissected cells were not labeled with alizarin, 2) they were located in the posterior part of cercarial body, 3) alongside with non-stained PA gland cells, alizarin-labeled CA gland cells were visible on the particular section of cercarial body to ensure that individual cercaria is stained properly. Nine biological replicates of both CA and PA glands, each containing 2.5 × 10^6^ μm^3^ of gland cell content, were microdissected. These were pooled in groups of three, resulting in 3 technical replicates of 7.5 × 10^6^ μm^3^ for each type of the glands.

### 2.11. Sample preparation for MS analysis

Three protocols were tested in order to optimize the process:

#### 2.11.1. In-solution digestion protocol

Microdissected cercarial tissues were resuspended in 30 μl of lysis buffer: 1% sodium deoxycholate (SDC, Merck), 10 mM Tris(2-carboxyethyl)phosphine (TCEP, Merck), 40 mM chloroacetamide (CAA, Merck), 100 mM triethylammonium bicarbonate (TEAB, ThermoScientific). Proteins were denatured, reduced, and alkylated in one step at 95 °C for 15 min and digested in lysis buffer using 0.5 μg of MS grade trypsin (ThermoScientific) per sample at 37 °C overnight. After digestion, the samples were acidified with trifluoroacetic acid (TFA, Merck) to 1% final concentration. SDC was removed by extraction to ethylacetate according to Masuda et al. (2008). Samples were desalted using in-house made Stage Tips filled with C18 resin (Empore) (Ishihama et al., 2006). Briefly, Stage Tips containing 3 layers of C18 extraction disks (Empore) were washed with a series of solutions as follows: 1) 50 μl of methanol 2) 50 μl of water, 90% acetonitrile (ACN, Merck), 0.1% TFA 3) 50 μl of water, 2% ACN, 0,1% TFA. After washing, samples were loaded, desalted using 100 μl of 2% ACN with 0.1% TFA, and peptides were eluted using 80 μl of 60% ACN with 0.1% TFA. Eluted peptides were dried and resuspend in 20 μl of loading solution (0.1% TFA, 2% ACN) prior to MS analysis.

#### 2.11.2. SP3 protocol

The protocol was adopted from Hughes et al. (2014). Microdissected cercarial tissues were resuspended in 20 μl of lysis buffer: 1% sodium dodecyl sulfate (SDS, Merck), 10 mM dithiothreitol (DTT, Merck), 50 mM 4-(2-hydroxyethyl)-1-piperazineethanesulfonic acid (HEPES, Merck), pH 8.5. The samples were transferred into 1.5 ml microtubes, 20 μl of 2,2,2-trifluoroethanol (TFE) was added, and sonication was performed by three short pulses using Sonoplus sonicator (Bandelin). Then, 0.75 μl of 0.1% formic acid (FA, ThermoScientific) was added. Microtubes were incubated at 95 °C for 5 min, placed for 30 s on ice, and then incubated at 45 °C for 30 min. The mixture was supplemented with 5 μl of 400 mM iodoacetamide (IAA, Merck) in 50 mM HEPES pH 8.5. Tubes were incubated at 24 °C in the dark for 30 min. To quench alkylation reaction, 5 μl of 200 mM DTT was added. Samples were dried using vacuum concentrator Modul 4080C (Hanil), resuspended in 8 μl of water, and sonicated for 5 min in a bath sonicator (Elma). Subsequently, 2 μl of bead-mix (SpeedBead Magnetic Modified Particles, GE Healthcare), 5 μl of 1% FA, and 15 μl of ACN were added to allow proteins to bind onto beads. After 10 min the supernatant was discarded using a magnetic rack and samples were washed with 2 × 200 μl of 70% ethanol and then with 180 μl of ACN. Dry beads were resuspended in 10 μl of 50 mM ammonium bicarbonate buffer (Merck) and 1 μg of trypsin and 0.1 μg of LysC were added. Samples were digested at 37 °C overnight, then desalted by washing beads with 2 × 200 μl of ACN in a magnetic rack. Beads were reconstituted using 9 μl of 2% dimethylsulfoxid (DMSO) in water and sonicated for 1 min in a bath sonicator (Elma). Finally, the beads were removed using a magnetic rack and 1 μl of 10% FA was added to the peptide solution.

#### 2.11.3. In-StageTips protocol

Microdissected cercarial tissues or particular penetration glands were processed by In-StageTips (IST) method according to Kulak et al. (2014). Briefly, pooled microdissects of particular gland cells were covered by 30 μl of lysis buffer (1% SDC, 10 mM TCEP, 40 mM CAA, 100 mM TEAB) and precisely pipetted from the adhesive caps into enclosed StageTips reactors containing styrenedivinylbenzene-reverse phase sulfonated (SDB-RPS) disks (Empore), under a constant stereomicroscopic supervision. Proteins were denatured, reduced, and alkylated in one step at 95 °C for 15 min. Proteins were digested using 0.5 μg of MS grade trypsin (ThermoScientific) per sample at 37 °C overnight. Digestion was stopped by 150 μl of 1% TFA. To remove SDC, reactors were washed twice with 150 μl of washing solution (50% ethyl acetate in 0.2% TFA). Samples were subsequently washed once with 100 ul of 0.2% TFA. Peptides were eluted with 2 × 50 μl of elution buffer (5% ammonium hydroxide, 80% ACN). Eluted peptides were dried and resolved in 20 μl of loading solution (0.1% TFA, 2% ACN, water).

### 2.12. Nanoflow liquid chromatography-tandem mass spectrometry

Nano Reversed phase column (EASY-Spray column, 50 cm × 75 µm ID, PepMap C18, 2 µm particles, 100 Å pore size) was used for LC/MS-MS analysis. Mobile phase A was 0.1% formic acid in water, mobile phase B was composed of 0.1% formic acid in ACN. Samples were loaded onto the trap column (Acclaim PepMap300, C18, 5 µm, 300 Å Wide Pore, 300 µm × 5 mm, 5 Cartridges) for 4 min at 15 μl/min. A loading buffer consisted of water, 2% ACN, and 0.1% TFA. Peptides were eluted with mobile phase B gradient from 4% to 35% B in 60 min. Eluting peptide cations were converted to gas-phase ions by electrospray ionization and analyzed on a Thermo Orbitrap Fusion device (Q-OT-qIT, Thermo). Survey scans of peptide precursors from 350 to 1400 m/z were performed at 120K resolution (at 200 m/z) with a 5 × 10^5^ ion count target. Tandem MS was performed by isolation at 1.5 Th with the quadrupole, HCD fragmentation with normalized collision energy of 30, and rapid scan MS analysis in the ion trap. The MS-MS ion count target was set to 10^4^ and the max injection time was 35 ms. Only those precursors with a charge state 2–6 were sampled for MS-MS. The dynamic exclusion duration was set to 45 s with a 10 ppm tolerance around the selected precursor and its isotopes. Monoisotopic precursor selection was turned on. The instrument was run in top speed mode with 2 s cycles (Hebert et al., 2014).

### 2.13. MS data analysis

Obtained raw data were analyzed and quantified with the MaxQuant software version 1.6.1.0. (Cox et al., 2014). The false discovery rate was set to 1 % for both proteins and peptides, and a minimum length of seven amino acids was specified. The Andromeda search engine was used for the MS-MS spectra search against the protein database (translated transcriptome) of cercariae and schistosomula of *T. szidati* (Leontovyč et al., 2019). In the case of microdissected penetration glands, the MS-MS spectra were additionally searched against protein databases (translated genomes) of other selected schistosome species *T. regenti* (trichobilharzia_regenti.PRJEB4662.WBPS14), *S. mansoni* (schistosoma_mansoni.PRJEA36577.WBPS14), and *S. japonicum* (schistosoma_japonicum .PRJEA34885.WBPS14), downloaded from the WormBase (https://parasite.wormbase.org). Enzyme specificity was set as C-terminal to arginine and lysine (variant C-terminal to lysine was also added in the case of SP3 method due to the LysC enzyme used), also allowing cleavage at proline bonds, and a maximum of two missed cleavages. Carbamidomethylation of cysteine was selected as fixed modification and N-terminal protein acetylation and methionine oxidation as variable modifications. The “match between runs” feature of MaxQuant was used to transfer identifications to other LC/MS-MS runs based on their masses and retention time (maximum deviation 0.7 min). Quantification was performed with the label-free algorithm described previously (Cox et al., 2014). After raw data evaluation in MaxQuant, further analysis was performed using Perseus software (Tyanova et al., 2016) versions 1.6.2.3 and 1.6.10.43. First, the data were transformed into binary logarithms. Reverse hits, hits identified only by site, and contaminants were filtered out. Student’s t-test performed in Perseus with permutation-based false discovery rate (FDR) calculation (250 randomizations) was used in the evaluation of protein spectra in experiments dealing with fixation and staining procedures (FDR = 0.05, S0 = 1), and also in the analysis of penetration gland contents (FDR = 0.05, S0 = 0.1). Proteins identified in all experiments were further annotated using non-redundant NCBI database (BLASTp; E-value cut-off 10^−5^). Proteins identified during penetration gland analyses were annotated as follows in order to fit the annotation closely to *T. szidati*. In case there were more protein identifiers matching a common set of peptides, only one protein identifier was selected for protein annotation in the following order: 1) according to *T. szidati*, 2) according to *T. regenti*, 3) according to *S. mansoni*, and 4) according to *S. japonicum*. Only the first protein identifier which corresponded to the selected species of parasite (as above) was chosen for protein annotation. Volcano plots were created using Instant Clue software version 0.5.3 (Nolte et al., 2018). In all experiments (except the evaluation of the fixation and staining procedures), the identified proteins were further annotated using Kyoto encyclopedia of genes and genomes (KEGG) (Kanehisa and Goto, 2000), (BLASTp; E-value cut-off: 10^−5^) and the protein identifications were further clustered into protein families according to KEGG annotation.

In the case of penetration gland content analysis, reliably identified proteins were further annotated using InterProScan search, and corresponding Gene Ontology (GO) IDs were obtained. After GO IDs merge in OmicsBox software (version 1.3.11; BioBam), GO enrichment analyses (Fisher’s exact test; filter mode P-value; Filter value: 0.05) regarding molecular functions were performed. Enzyme code distribution (main classes and the abundant subclasses) was performed in OmicsBox. In addition, blast against peptidase units and inhibitor units in MEROPS database (version 12.1) (E-value: 10^−5^) was performed.

### 2.14. Proteome vs. transcriptome: quantification of peptidases and peptidase inhibitors

Peptidases and inhibitors playing a possible role in the invasion of *T. szidati* cercariae into the host were selected and quantified using their measured proteomic and transcriptomic values. Protein intensities from the current proteomic study and measured values of transcripts per million (TPM) from the transcriptomic study (Leontovyč et al., 2019) were employed for comparison.

### 2.15. Artificial assembly of the total proteome of *T. szidati* cercarial bodies

Taking into account the effect of alizarin staining on proteomic analyses, the total proteome was compiled based on data obtained in two separate experiments, namely a) analysis of penetration gland content (described in 2.8), and b) from raw data of both unstained/unfixed and alizarin-stained cercarial bodies (without tails) (see 2.10 staining experiment). Using MaxQuant, selected raw data from the staining experiment were re-searched against protein databases of *T. szidati, T. regent*i, *S. mansoni*, and *S. japonicum*. Only reliably identified proteins present in at least three of the five replicates in at least one body sample type (unstained/unfixed or stained with alizarin), and those present in at least two of the three replicates in at least one type of a gland sample (CAg or PAg) were used to assemble the total proteome.

## 3. Results

### 3.1. Optimization of the amount of material and the method of sample preparation for MS analysis

In-solution digestion (ISD), SP3 and In-StageTips (IST) protocols were applied for various volumes of microdissected cercarial tissues and the results of following MS analyses were compared. Only protein identifications with valid MS precursor intensity values were included. When comparing the number of proteins captured by individual methods, both IST and ISD methods provided better results than the SP3 protocol over almost whole range of sample volumes (Fig. 1A, Supplementary Table S1 sheets A – O, and Supplementary Fig. S2). The IST protocol was proven to be the best one for most KEGG protein categories (Fig. 1B). Moreover, this protocol requires fewer steps in terms of material transfer, which may be critical in experiments with hard-to-reach samples of minimal volumes, and can also be done in the shortest time. Therefore we decided to use the IST method in all following experiments. Further testing of the IST method with an extended range of sample volumes led us to a decision to apply the volumes of 7.5 × 10^6^ μm^3^ of the microdissected material in all following analyses (Fig. 1C and Supplementary Table S3 sheets A – F). This proved to be optimal in terms of effectiveness considering the sample gain/processing time and the number of identified proteins.

**Fig. 1.**
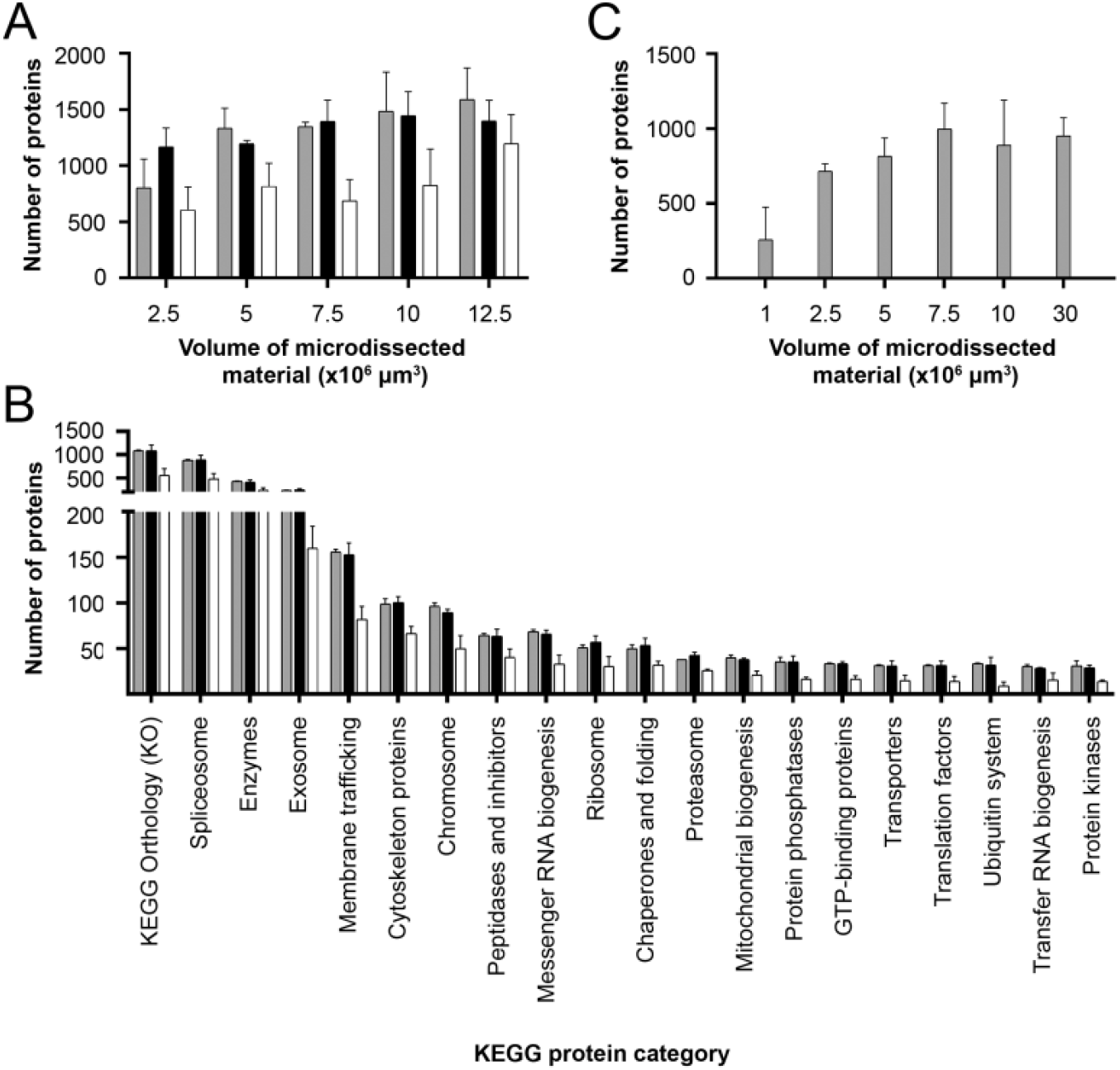
Selection of the optimal sample volume and the method of sample processing for MS analysis of microdissected cercarial tissues. **(A)** Comparison of In-StageTips (IST, grey), In-solution digestion (ISD, black) and “SP3” (SP3, white) protocols for processing of microdissected material for MS analysis based on numbers of MS-identified proteins. (**B)** Comparison of the numbers of MS-identified proteins in samples processed by ISD, IST or SP3 method. Proteins were categorized according to KEGG (Kyoto encyclopedia of genes and genomes). The results for the microdissected tissue volume of 7.5 × 10^6^ μm^3^ are presented in this graph (only the 20 most abundant protein categories are shown). Comparisons based on the other volumes tested has shown similar trends with the exception of the volume 2.5 × 10^6^ μm^3^ (also see Supplemetary Fig. S2). **(C)** A test of the IST method using an extended range of tissue volumes. The volume of 7.5 × 10^6^ μm^3^ of the microdissected material processed by the IST method provided optimal results and was used in all following analyses. Only protein identifications with valid MS precursor intensity values were used to generate graphs in GraphPad Prism (version 8.0.1). Results in graphs are presented as mean values ± SD.

### 3.2. Optimization of tissue preservation protocol

Dorsal and lateral views of a whole living intact cercaria of *T. szidati* with description of parts of cercarial body are shown in Fig. 2A and Fig. 2B, respectively. The microscopic evaluation of the appearance of sectioned cercarial body tissues in non-fixed (NF), paraformaldehyde-(PF) or ethanol-treated (EtOH) samples revealed a well-maintained structure in NF samples, with only minor cracks present (Fig. 2C). In EtOH-fixed samples, the structure was relatively well preserved, but moderate body shrinkage occurred (Fig. 2D). In PF-fixed samples, excessive body shrinkage was obvious which hampered gland definition and, moreover, has led to significant loss of penetration gland contents due to their release during fixation (Fig. 2E).

**Fig. 2.**
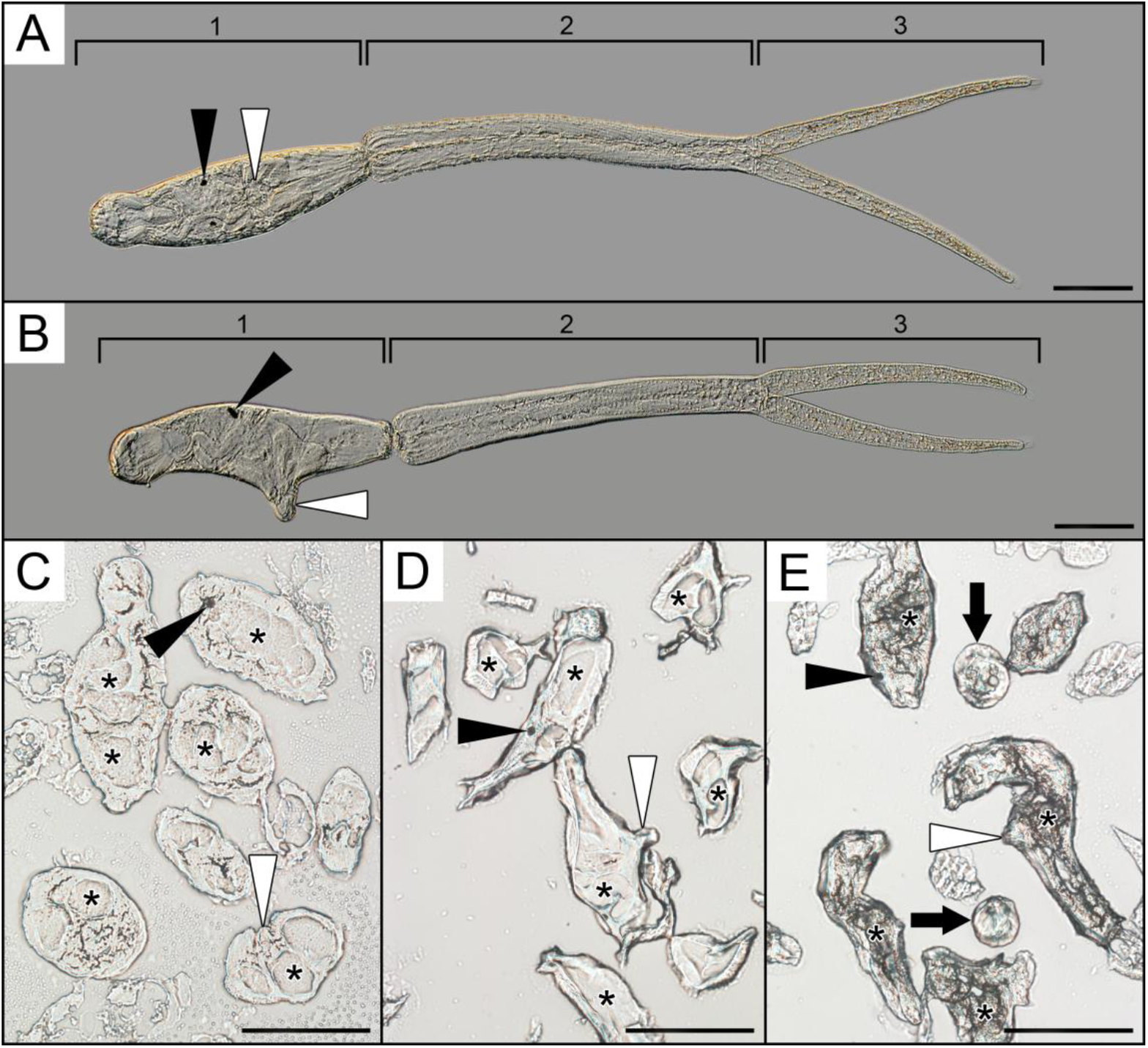
Examples of a living intact cercaria and fixative-preserved cryosectioned cercariae. **(A)** Living non-fixed cercaria of *Trichobilharzia szidati* in dorsoventral and **(B)** lateral orientation under Nomarsky interferrence contrast. Region 1 - cercarial body (head), region 2 - cercarial tail stem, region 3 - cercarial tail furca. **(C – E)** Cryosections on microscopic slides as observed during laser microdissection. **(C)** Well-preserved structure of cercariae in non-fixed samples, only minor cracks are present. Glands are apparently visible. **(D)** Ethanol-fixed samples with moderate body shrinkage. **(E)** Samples fixed in depolymerized paraformaldehyde with excessive body shrinkage. Position of the glands is unclear. Bright-field. Black asterisks indicate some of individual gland cells in cercarial bodies. Black arrows show cercarial tails. Black arrowheads point to pigmented eye spots, white arrowheads point to ventral sucker (acetabulum). Scale bar = 100 µm. Photographs were taken under Olympus BX51 microscope equipped with Olympus DP72 camera.

The numbers of proteins detected by MS analysis in microdissected samples after different types of treatment were compared. A cumulative number of identified individual proteins reached 1636. Only those detected in at least three out of six replicates of at least one type of tissue preservation were considered as reliably confirmed. After setting this requirement, the cumulative number of reliably identified proteins dropped to 781. Protein numbers varied just slightly among different types of sample treatment (NF 770, EtOH 746, and PF 758) (Fig. 3A). Using label-free quantification, the MS results of different types of tissue processing were compared each other. The abundance of most of the proteins did not differ significantly between particular pairs of samples represented by different types of tissue processing (Fig. 3B – D). A detailed list of identified proteins can be viewed in Supplementary Table S4 that is complemented with Supplementary Fig. S5.

**Fig. 3.**
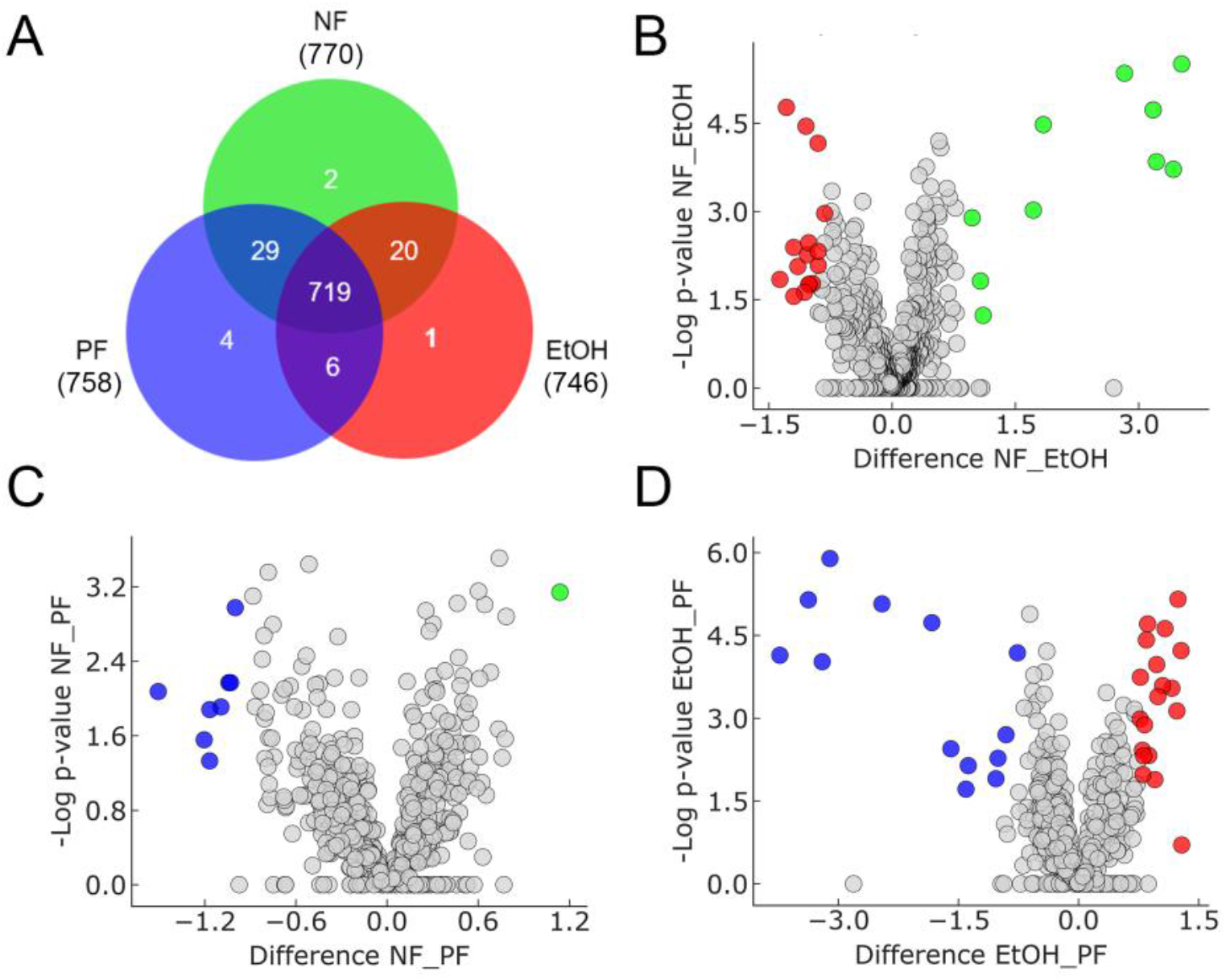
Comparison of numbers and abundance of reliably MS-identified proteins in cryosections of cercarial bodies after three types of tissue treatment. **(A)** The Venn diagram shows the numbers of reliably identified proteins in non-fixed (NF, green), ethanol-fixed (EtOH, red), and paraformaldehyde-fixed (PF, blue) cercarial tissues. Volcano plots **(B – D)** show differences in protein abundances between two particular types of tissue processing. Dots represent individual proteins; those being significantly more abundant are marked in colour as: green = NF samples, red = EtOH samples, blue = PF samples. **(B)** Comparison of NF and EtOH samples. **(C)** Comparison of NF and PF samples. **(D)** Comparison of EtOH and PF samples. Label-free quantification was applied (Student’s t-test, FDR = 0.05, S0 = 1).

### 3.3. Optimization of staining protocol for cercarial tissues and penetration glands

Toluidine blue did not stain any type of the glands specifically, while lithium carmine stained specifically the PAg. In our experimental setting, trypan blue stained the PAg, but also induced release of (mixed) gland products which adhered to cercarial bodies. Therefore it was not possible to use it. Alizarin stained specifically the CAg, however, staining was only possible in the stage of still living cercariae just before freezing and cryosectioning (Fig. 4).

**Fig.4.**
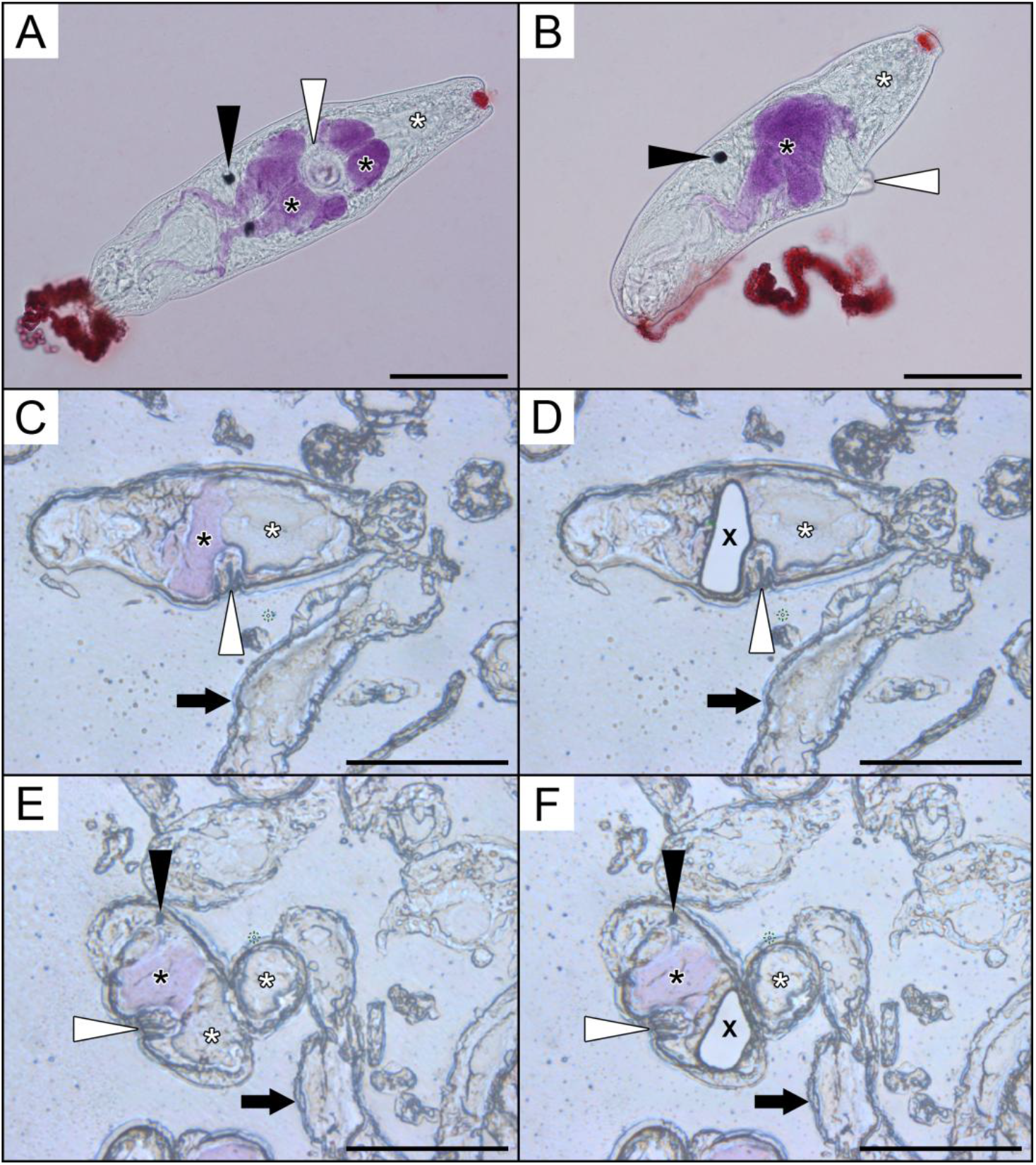
Circumacetabular penetration glands of *Trichobilharzia szidati* cercariae stained by alizarin. **(A)** Dorsoventral and **(B)** lateral view of intravitaly stained cercarial body with pale violet coloration in circumacetabular gland cells and their ducts. **(C - F)** The process of microdissection (prior and post dissection) on cryosectioned cercariae stained with alizarin for circumacetabular **(C - D)** and postacetabular **(E - F)** glands. Black asterisks indicate selected individual circumacetabular gland cells, white asterisks show position of postacetabular glands. X indicates the empty area post dissection. Black arrows show cercarial tails without glands. Black arrowheads point to pigmented eye spots, white arrowheads point to ventral sucker (acetabulum). Scale bar = 100 µm.

The number of proteins detected by MS analysis in microdissected samples of cercarial bodies after application of different histological dyes was compared. A cumulative number of identified individual proteins reached 1301. Only those detected in at least three out of five replicates of at least one type of staining were considered as reliably confirmed. After setting this requirement, the cumulative number of reliably identified proteins dropped to 962 (936 in unstained samples, 582 in toluidine blue, 956 in alizarin, 909 in trypan blue, and 789 in lithium carmine). The comparison between stained and unstained samples using label-free quantification showed significant differences in protein levels (Fig. 5). A detailed list of identified proteins can be viewed in Supplementary Table S6 that is complemented with Supplementary Fig. S7. Regarding the number of proteins obtained, alizarin and trypan blue gave the best results which were comparable with those from unstained samples. In addition, alizarin showed the slightest differences in protein abundances when compared with unstained samples (Table 1).

**Table 1.** Comparison of numbers of reliably MS-identified proteins in unstained and stained cryosections of cercarial bodies.

| Histological dye | Identified proteins | Exclusive proteins | More abundant in unstained | More abundant in used dye |
| --- | --- | --- | --- | --- |
| toluidine blue | 582 | 0 | 182 | 140 |
| alizarin | 956 | 4 | 51 | 69 |
| trypan blue | 909 | 0 | 133 | 106 |
| lithium carmine | 789 | 0 | 162 | 67 |
| unstained | 936 | 1 | N/A | N/A |

**Fig. 5.**
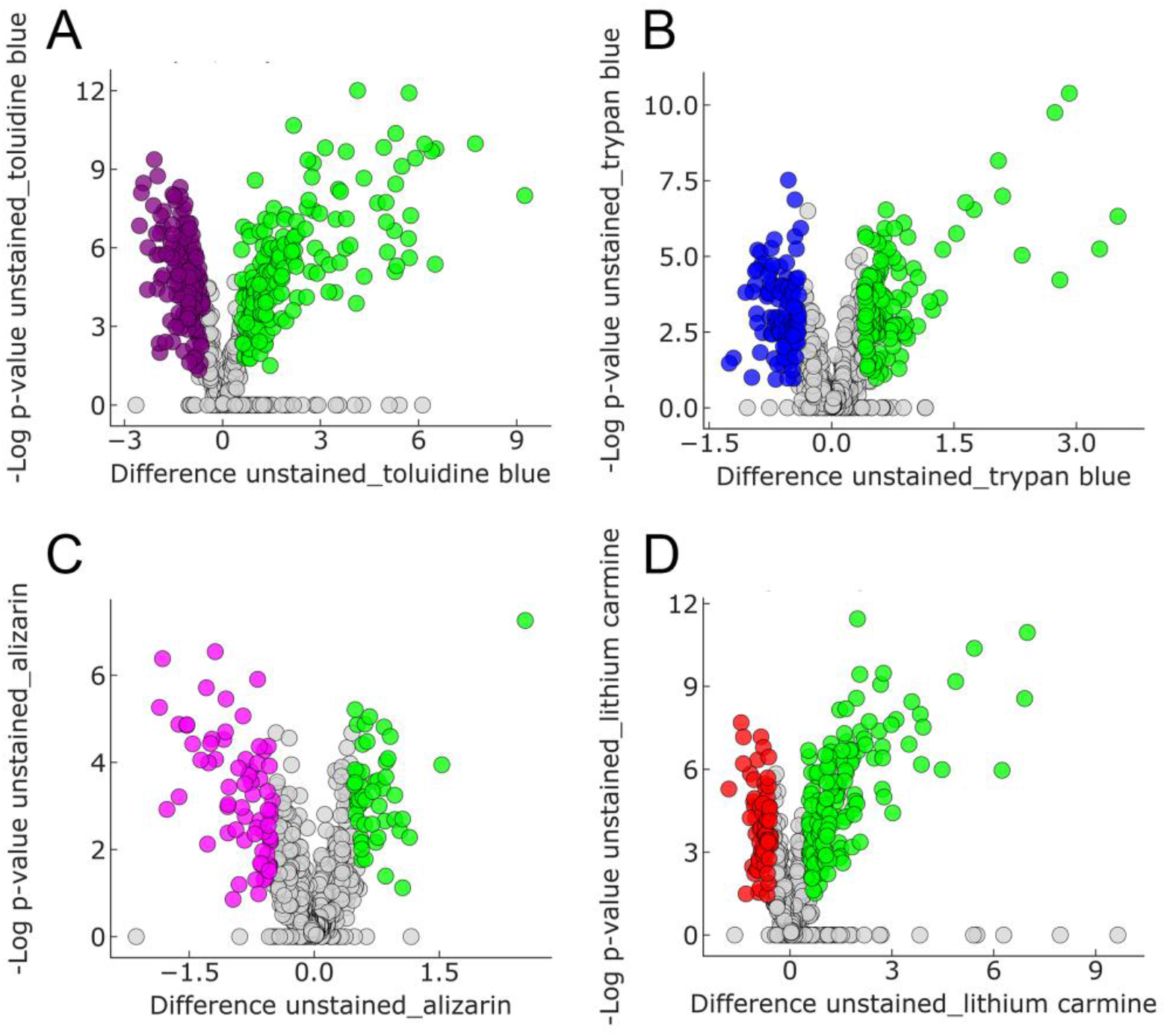
Comparison of the abundance of reliably MS-identified proteins in unstained and stained cryosections of cercarial bodies. Comparison of stained and unstained samples using label-free quantification (Student’s t-test with FDR = 0.05, S0 = 1) showed significant differences in protein abundances. Volcano plots **(A – D)** show differences in protein abundances between an unstained sample and a sample stained with a particular dye. Dots represent individual proteins; those being significantly more abundant are marked in colour as: green = unstained sample, purple = toluidine blue, pink = alizarin, blue = trypan blue, red = lithium carmine. Specific numbers of the identified proteins are shown in Table 1. **(A)** Comparison of unstained vs. toluidine blue-stained samples. **(B)** unstained vs. trypan blue. **(C)** unstained vs. alizarin. **(D)** unstained vs. lithium carmine. Alizarin staining was applied on living cercariae prior cryopreservation, other dyes (toluidine blue, lithium carmine, and trypan blue, were applied just to cryosections of cercariae.

### 3.4. The effect of processing time on sample stability at RT

In an attempt to check protein stability in non-fixed microdissected samples of cercarial body tissues, and also to simulate prolonged microdissection process, samples of equal volumes were exposed to RT for various time periods. Only protein identifications with valid MS precursor intensity values were used for comparison. In general, the numbers of MS-identified proteins did not differ considerably among different times (Fig. 6). A detailed list of identified proteins for each time interval can be viewed in Supplementary Table S8 sheets A - F. By comparing the spectra of proteins categorized according to KEGG, the numbers of proteins in individual categories did not differ significantly among the time intervals tested (see Supplementary Table S9). The exposure of the non-fixed samples to RT up to 24 h had obviously no considerable effect on the overall stability of proteins.

**Fig. 6.**
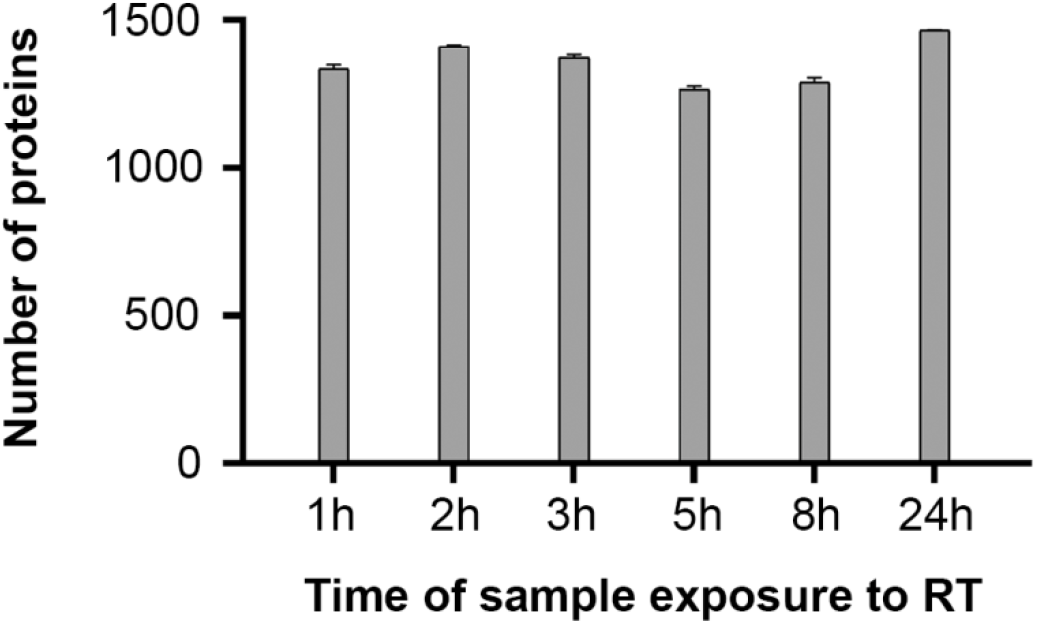
The effect of time and room temperature on numbers of MS-identified proteins in microdissected non-fixed cercarial tissues. Numbers of proteins for each time interval are shown, the time of exposure to RT includes the time needed for the microdissection process. Only protein identifications with valid MS precursor intensity values were used to create the graph in GraphPad Prism (version 8.0.1). Results are presented as mean values ± SD.

### 3.5. Cercarial penetration gland analysis

#### 3.5.1. Identification of postacetabular and circumacetabular gland proteins and their classification

After separate microdissection of CAg and PAg (Fig. 4), their protein contents were analyzed. Using MS/MS-based proteomic analysis, we identified 3390 peptides (including 43 identified only by site) (Supplementary Table S10). From these, 3347 peptides were assigned to 792 proteins. Those detected in at least two out of three samples from either gland type were considered as reliably identified. After this requirement was set, the resulting 461 proteins (PAg 455 in total, 40 exclusives; CAg 421 in total, 6 exclusives; see Venn diagram in Fig. 7 and Supplementary Table S11, sheets A – B) were annotated according to NCBI, KEGG and MEROPS databases and classified by use of Gene Ontology. Using label-free quantification, 60 out of 415 common proteins differed significantly in abundance between CAg and PAg (48 were more abundant in PAg and 12 in CAg) (Fig. 7 and Supplementary Table S11, sheets C – D). Other 355 proteins were detected in both types of the glands but the differences in their abundance between particular glands were below the level of significance (Supplementary Table S11, sheet E). Annotated spectra for protein identifications based on 1 peptide are presented in Supplementary Fig. S12 that complements Supplementary Table S11. The 331 proteins which did not meet our criteria of reliable identification are listed in Supplementary Table S13.

**Fig. 7.**
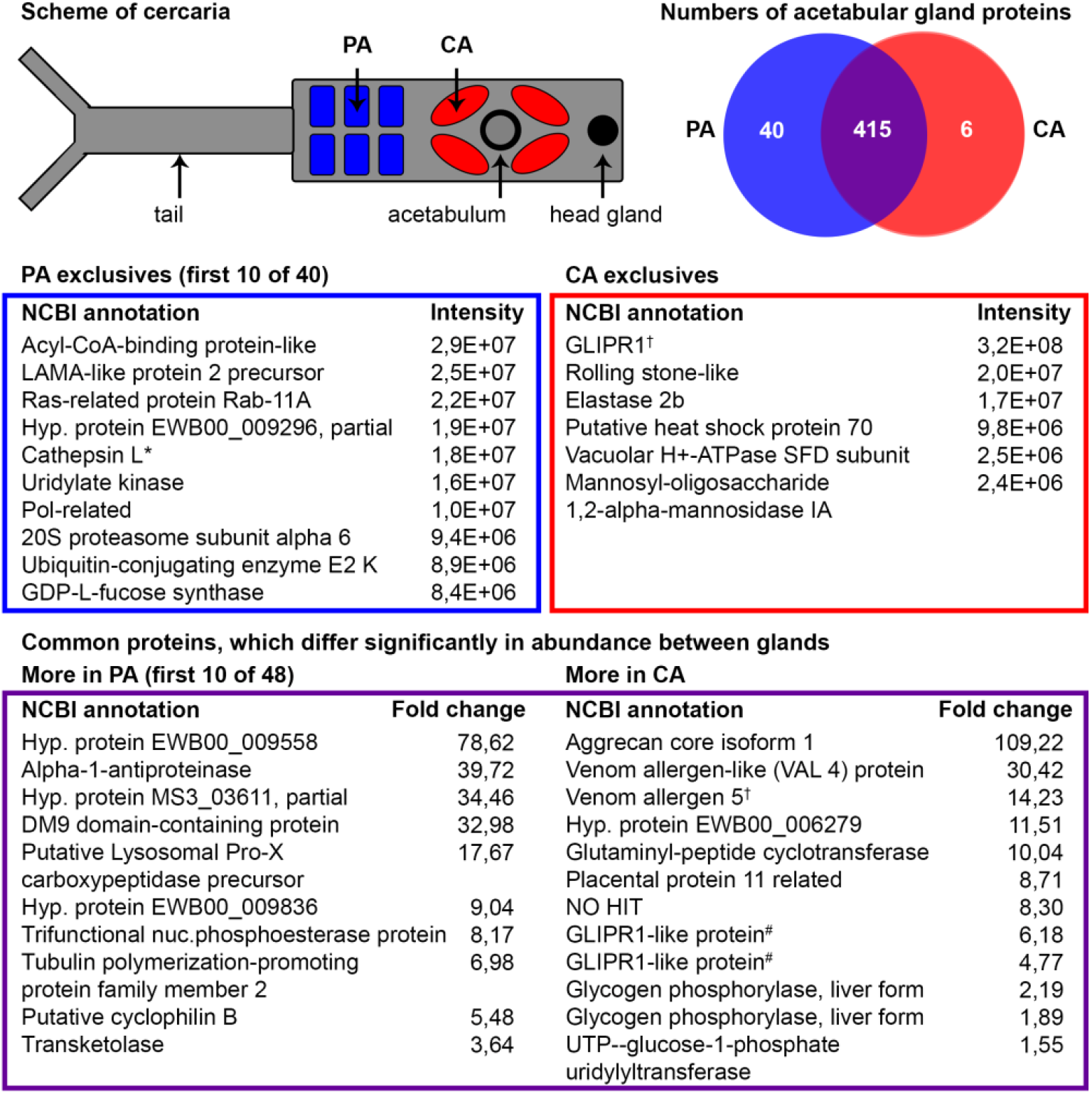
Distribution of exclusive and most abundant proteins between acetabular glands of *Trichobilharzia szidati* cercariae. By comparing the contents of circumacetabular (CA) and postacetabular (PA) glands, it was possible to reliably identify 461 proteins in total, where 40 were exclusively present in PA glands and 6 exclusively in CA glands. Using label free quantification (Students t-test with FDR = 0.05, S0 = 0.1), levels of 60 proteins differ significantly between PA and CA glands (40 were more abundant in PA glands, 12 more abundant in CA glands). A detailed list of identified proteins is a part of Supplementary Table S11. *SmCL3-like; ^†^an ortholog of SmVAL18; ^#^an ortholog of SmVAL10.

Reliably identified proteins were further clustered into protein families according to KEGG (Supplementary Table S11, sheet F). The highest numbers of proteins in both types of the glands were associated with the enzymes and exosome categories. Other widely represented proteins included mainly cell house-keeping proteins belonging to categories such as cytoskeleton, membrane trafficking, chromosome and associated proteins, chaperones and folding catalysts, mRNA biogenesis, GTP-binding, and mitochondrial biogenesis. Most were shared by both gland types and just a relatively small proportion of them were identified as specific for a particular gland type. However, among 182 KEGG-annotated enzymes, 29 % were exclusive or significantly more abundant in PAg while only 5 % were exclusive or more abundant in CAg. The distribution of enzyme classes (EC) in the glands according to Gene Ontology annotation is shown in Fig. 8. The most represented groups comprised hydrolases and transferases. Among hydrolases, those acting on peptide bonds (peptidases) and on acid anhydrides (GTPases, ATPases) were dominant. Noteworthy is also the occurrence of glycosylases of which some are exclusive for particular gland types (Fig. 9). The largest proportion of transferases in the glands included phosphotransferases, glycosyltransferases (especially in CAg), and acyltransferases (Fig. 10).

**Fig. 8.**
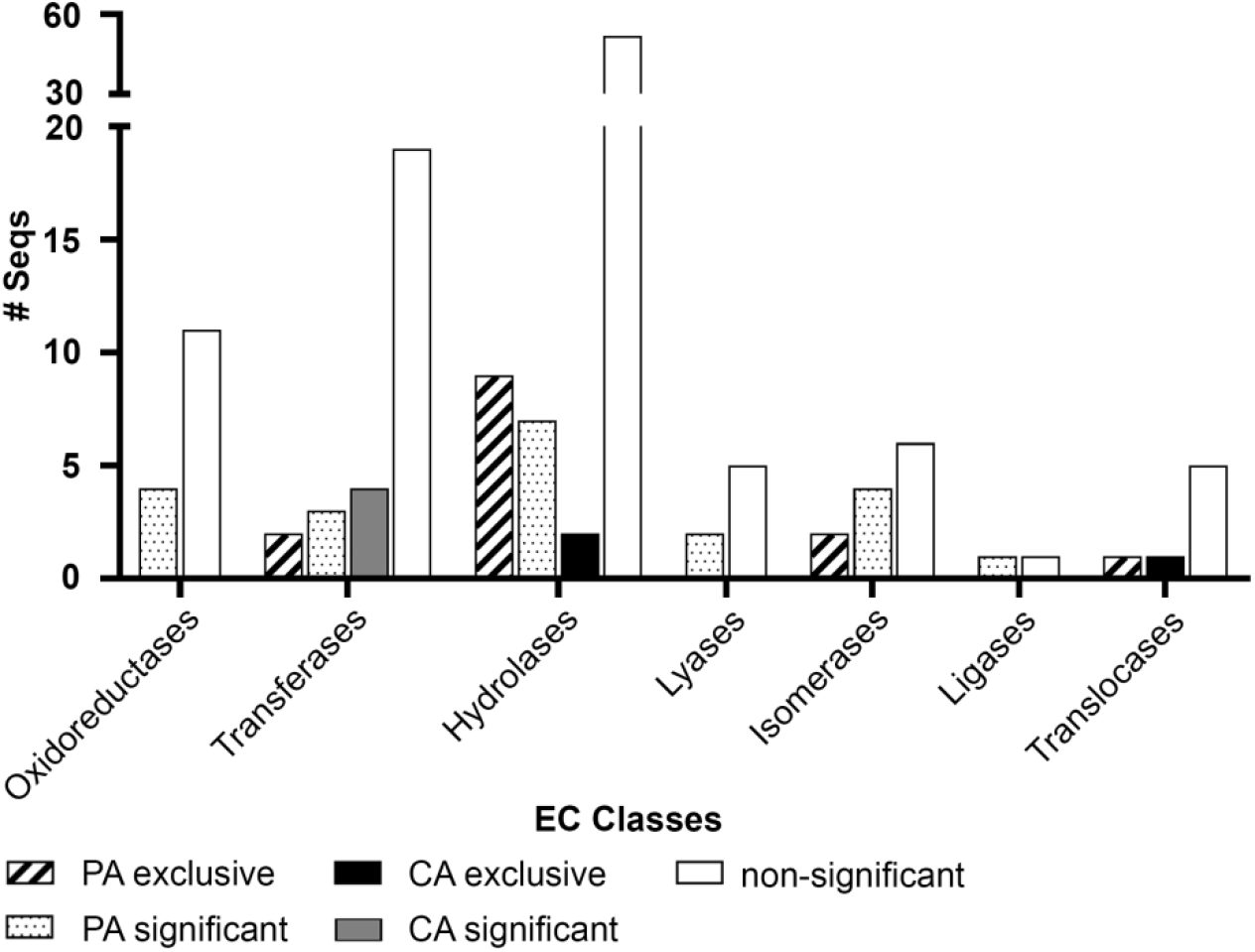
Distribution of enzyme main classes in the acetabular glands of *Trichobilharzia szidati* cercariae. The most represented groups comprised hydrolases and transferases. The distribution was performed in OmicsBox software (version 1.3.11).

**Fig. 9.**
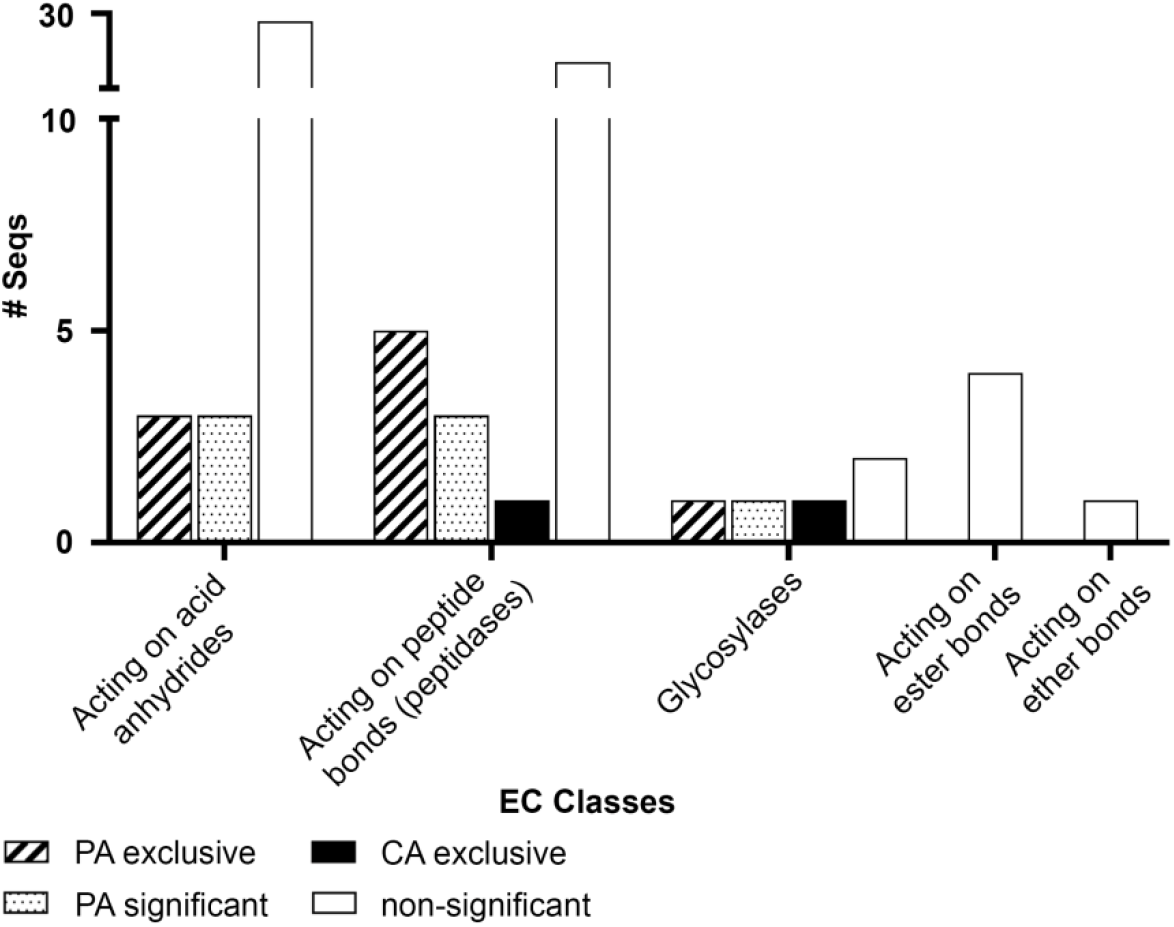
Distribution of hydrolases in the acetabular glands of *Trichobilharzia szidati* cercariae. The dominant groups were those containing, e.g., peptidases and GTPases or ATPases. Some glycosylases were exclusive for particular gland types. The distribution was performed in OmicsBox software (version 1.3.11).

**Fig. 10.**
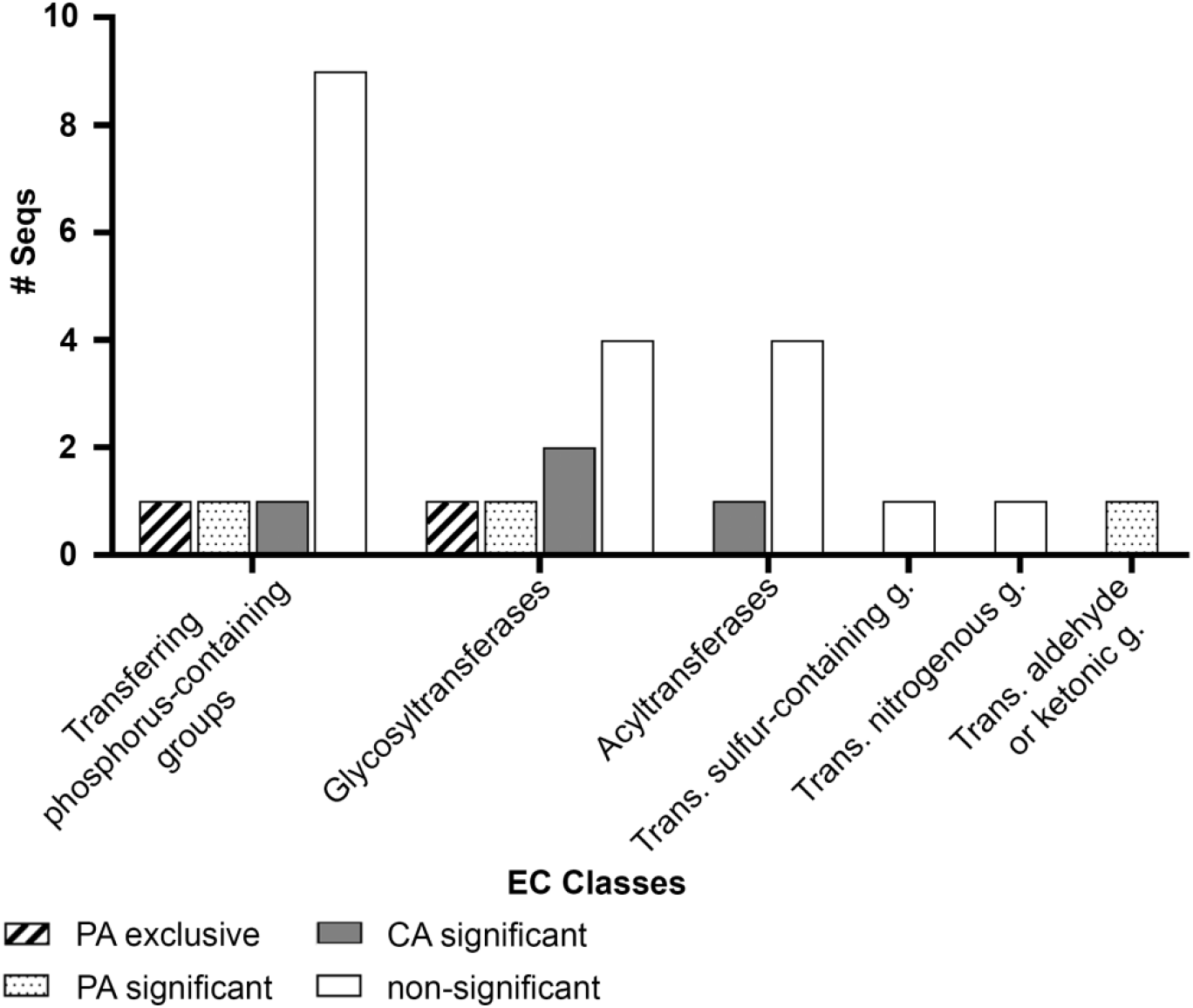
Distribution of transferases in the acetabular glands of *Trichobilharzia szidati* cercariae. Phosphotransferases, glycosyltransferases, and acyltransferases were the most prevalent groups between the two types of glands. The distribution was performed in OmicsBox software (version 1.3.11).

A Gene Ontology (GO) enrichment analysis regarding molecular functions has shown over-representation of GO terms related to catalytic activity (e.g. cysteine type peptidase activity or ubiquitinyl hydrolase activity) among PAg-exclusive proteins (Fig. 11A). In the case of PAg-significant proteins, GO terms related to the structural constituents of the cytoskeleton and also to catalytic activity (peptidyl-prolyl isomerase activity) were enriched (Fig. 11B). In the case of CAg-exclusive proteins, transporter activity and catalytic activity (proton-transporting ATPase, mannosidase and serine-type endopeptidase activity) were enriched (Fig. 12A). For CAg-significant ones, binding (pyridoxal phosphate binding) and catalytic activity-related GO terms were enriched (glycogen phosphorylase, glutaminyl-peptide cyclotransferase, UTP:glucose-1-phosphate uridylyltransferase, and endoribonuclease activity) (Fig. 12B).

**Fig. 11.**
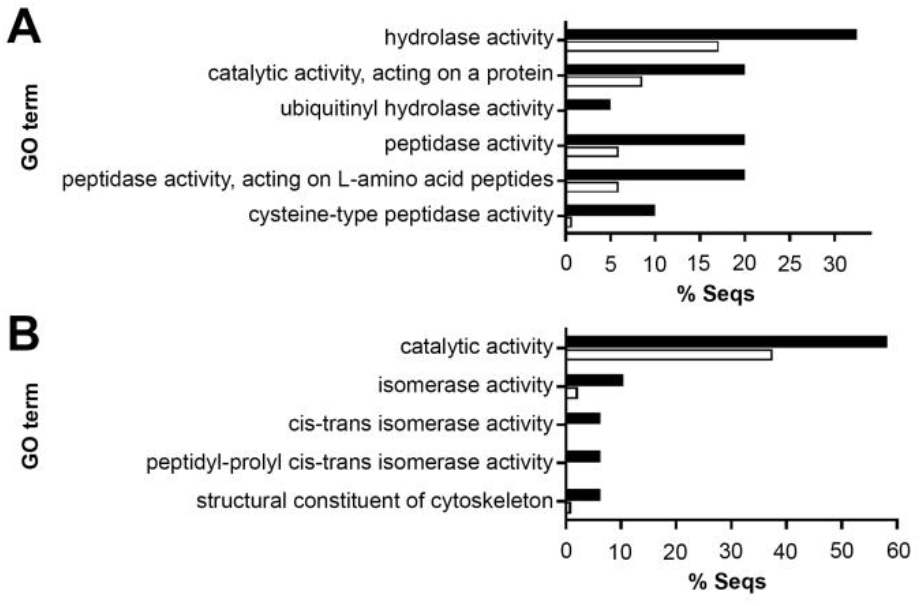
GO enrichment analysis: postacetabular glands (PAg) of *Trichobilharzia szidati* cercariae. GO enriched bar charts regarding molecular functions showing over-represented GO terms (Fisher’s exact test performed in OmicsBox software (version 1.3.11); filter mode P-value; Filter value: 0.05). Test file: black bars; reference file: white bars. **(A)** Among proteins exclusively identified in PAg (30 out of 40 proteins with GO terms), GO terms related to, e.g., cysteine type peptidase activity or ubiquitinyl hydrolase activity were enriched. **(B)** Among proteins significantly more abundant in PA glands (35 out of 40 proteins with GO terms), GO terms related to the structural constituents of the cytoskeleton and peptidyl-prolyl isomerase activity were enriched.

**Fig. 12.**
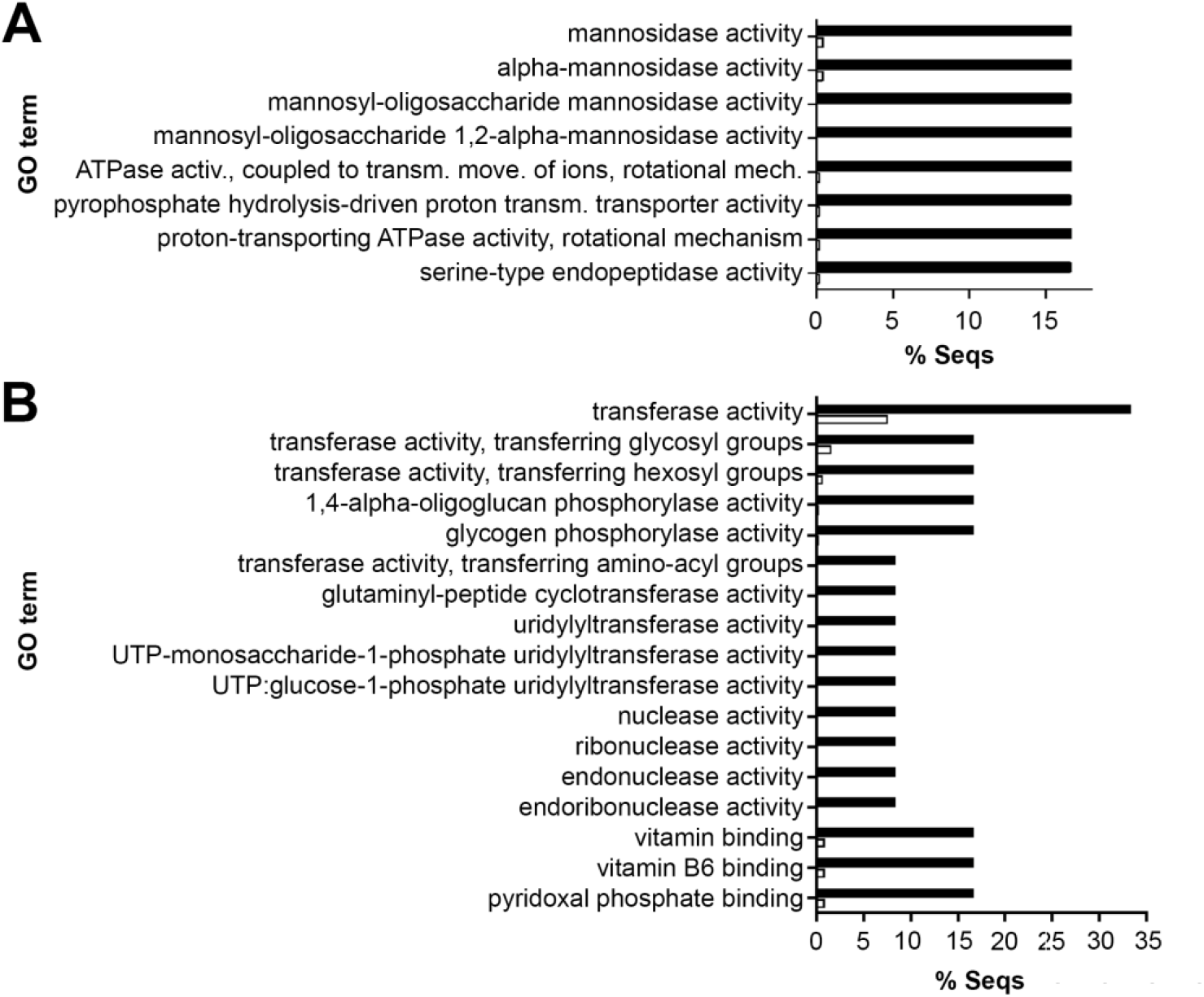
GO enrichment analysis: circumacetabular glands (CAg) of *Trichobilharzia szidati* cercariae. GO enriched bar charts regarding molecular functions showing over-represented GO terms (Fisher’s exact test performed in OmicsBox software (version 1.3.11); filter mode P-value; Filter value: 0.05). Test file: black bars; reference file: white bars. **(A)** Among proteins exclusively identified in CAg (3 out of 6 proteins with GO terms), GO terms related to proton-transporting ATPase, mannosidase and serine-type endopeptidase activity were enriched. **(B)** Among proteins significantly more abundant in CA glands (5 out of 12 proteins with GO terms), GO terms related to pyridoxal phosphate binding and glycogen phosphorylase, glutaminyl-peptide cyclotransferase, UTP:glucose-1-phosphate uridylyltransferase, and endoribonuclease activity were enriched.

#### 3.5.2. Peptidases and peptidase inhibitors

Based on the combination of NCBI, MEROPS, and KEGG annotations, 39 peptidases and 3 peptidase inhibitors were identified among the reliably identified proteins in the acetabular glands (Supplementary table S11, sheet G). There were differences in the annotations from different databases in cases of some proteins, therefore we decided to present here the consensual results. More than 1/3 of the peptidases are constituents of the proteasome assembly. All gland peptidases were assigned to appropriate catalytic types: 7 cysteine peptidases, 13 threonine peptidases, 13 metallo peptidases, 5 serine peptidases, and 1 aspartic peptidase.

Reliably identified peptidases (including proteasome subunits) accounted for roughly 8 % of reliably identified PAg proteins and 6 % of CAg proteins. Elastase 2b (according to NCBI annotation) was identified exclusively in the CAg as the third most abundant protein (Fig. 7). Exclusive for PAg were 4 cysteine peptidases (including SmCL3-like cathepsin L, being the fifth most abundant protein, Fig. 7), 2 threonine peptidases, 2 metallo peptidases, and 1 serine peptidase. Significantly higher abundance in PAg was observed for 2 serine, 2 metallo and 1 threonine peptidase. Represented in both types of glands, but without a significant differences in abundance, the following were identified: 10 threonine peptidases, 9 metallo peptidases, 3 cysteine peptidases including cathepsin B2 (this was just below the significance for greater representation in PAg), 1 aspartic peptidase, and 1 serine peptidase (Supplementary table S11, sheet G). Besides, among the proteins identified just in one of three replicates for each gland type, we found (according to NCBI annotation) an M17-family leucyl aminopeptidase being exclusive for PAg (Supplemetary table S13, sheet A). A family C19 unassigned peptidase, M08 family invadolysin, M01 family alanyl aminopeptidase, and C01 family dipeptidyl peptidase cathepsin C were exclusive for CAg (Supplementary table S13, sheet B). Another invadolysin, leishmanolysin and a M16 family metallopeptidase were common for both gland types (Supplementary table S13, sheet C).

As for peptidase inhibitors, annotations revealed 3 candidates (Supplementary table S11, sheet G). A serpin (Alpha-1-antiproteinase according to NCBI) was found at significantly higher levels in PAg (39 fold change, Fig. 7 and Supplementary table S11, sheet G). Two cystatins were found in both gland types with no significant difference. Additionally, peptidase inhibitors identified just in one of three replicates for each gland type included another serpin being exclusively in PAg (Supplemetary table S13, sheet A), and one more serpin superfamily member (alpha-1-antiproteinase 2) plus a potential Kunitz-type inhibitor exclusive for CAg (Supplemetary table S13, sheet B). Annotated spectra for the above-mentioned protein identifications based on just 1 peptide and present in just one of the three replicates for each gland type are presented in Supplementary Fig. S14 that complements Supplementary Table S13.

Comparison of the abundances of reliably identified non-proteasome peptidases and of peptidase inhibitors in *T. szidati* cercarial glands is shown in Fig. 13A. The abundances of their coding transcripts based on transcriptomic data from cercariae (Leontovyč et al., 2019) are in Fig. 13B. Measured values are presented in Supplementary Tables S11, sheet H.

**Fig. 13.**
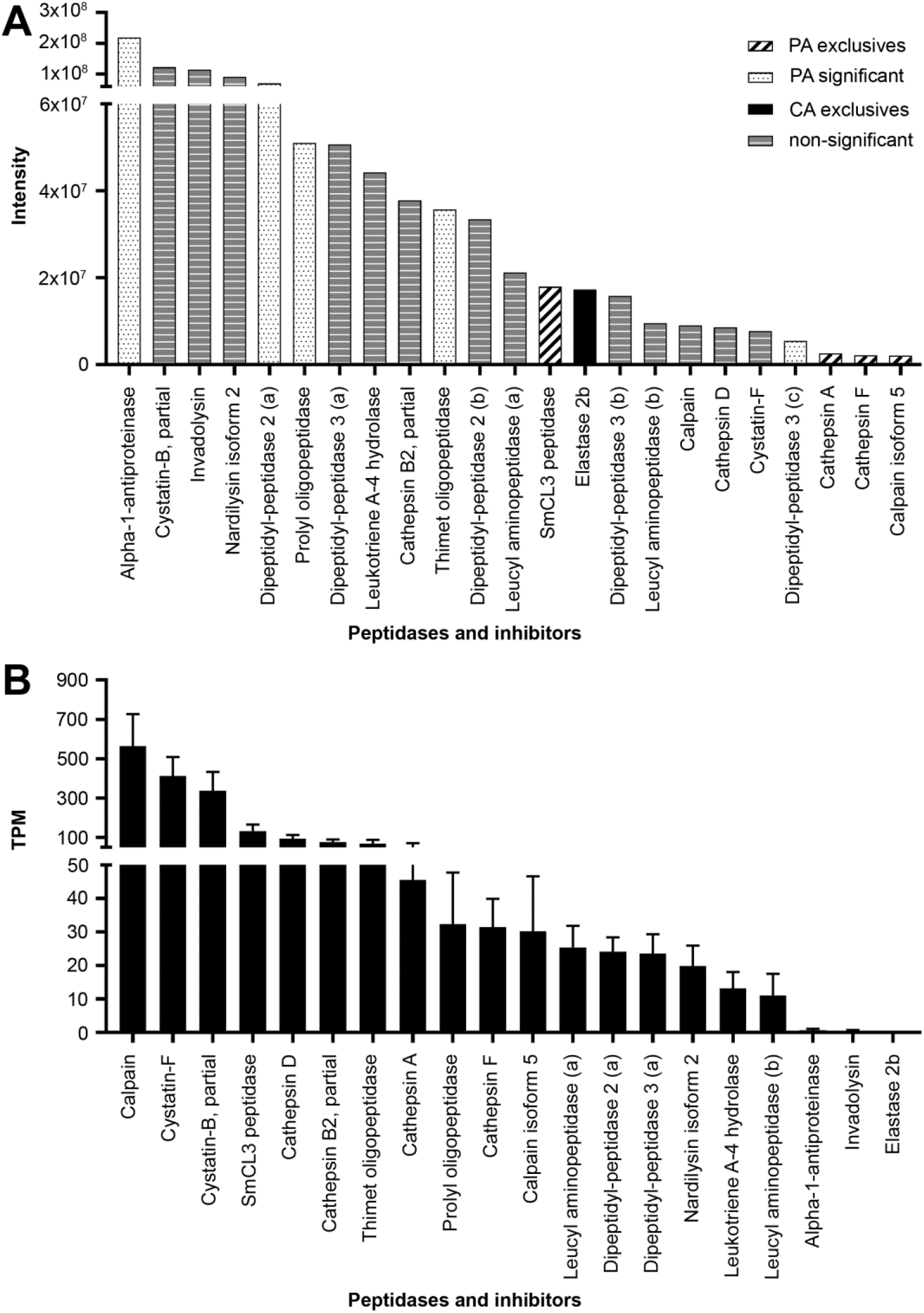
Peptidases and inhibitors playing a possible role in the invasion of *Trichobilharzia szidati* cercariae into host: comparison of their relative abundance in proteomic and transcriptomic assays. The graphs show a different situation regarding mRNA and proteins themselves. In some cases (alpha-1-antiproteinase, invadolysin, and elastase 2b) almost no transcripts were detected in contrast to proteomic study. These enzymes are probably not synthesized in cercariae. In contrast, the situation, e.g. with calpain is different. These results may help answering question concerning proteosynthesis in free-swimming cercariae. **(A)** Protein intensities from the current proteomic study. **(B)** The abundances of coding transcripts: measured values of transcripts per million (TPM).

#### 3.5.3. Non-peptidase proteins with putative roles in invasion of the host

Besides peptidases, several other proteins possibly involved in the invasion of the host skin by cercariae and in the establishment of the infection were also identified. These include, e.g., a higly abundant postacetabular LAMA-like protein containig phospholipase B domain, putative beta-thymosin, DM9 domain-containing protein, superoxide dismutase, glutathione-S-transferase etc. Conspicuous was the high abundance of cysteine-rich secretory proteins of the SCP/TAP family (Pfam accession number PF00188) in the CAg, known also from other trematodes and nematodes. These included a CA-exclusive GLIPR1 (VAL18) protein and CA-significant VAL4 and VAL5 proteins. Other orthologs of VAL proteins (GLIPR1-like proteins) were localized in both types of acetabular glands. Another relatively abundant and important group of proteins found in *T. szidati* acetabular gland proteomes, much like in mixed gland ESPs of other schistosomes, were molecular chaperones, heat shock proteins (HSPs). A HSP70 family member was identified exclusively in the CAg and some other homologs occurred among the common gland proteins. Other HSPs (60, 20, 10) were also present among common proteins.

Similarly to other schistosomes, we also identified a considerable number of different calcium-binding or regulatory proteins in both gland types, namely PAg-significant calcium-binding EF-hand-containing glutathione-S-transferase, and Ca^2+^-dependent proteins common to both gland types such as calcium-binding EF-hand domain-containing acyl carrier protein, kinases, myosin light chain, calmodulins, calreticulin, calbindin-32, calpains, calponins, annexins, and others (Supplementary Tables S11, sheets A – E).

### 3.6. Total cercarial body proteome

Although this paper was focused on characterization of differential proteomes of cercarial penetration glands, we obtained a lot of information about the proteome of the cercarial bodies (heads) exluding tails from numerous replicates used for optimization of proteomic methods. Therefore we present the data as a by-product of the study (Supplementary table S15). Taking into account the effect of alizarin staining, the proteome was artificially assembled from reliably identified proteins from two separate experiments a) analysis of penetration gland content, and b) from raw data of unstained/unfixed and alizarin-stained samples of microdissected areas of cercarial bodies excluding tails (staining experiment, re-evaluated). Based on NCBI annotation, we present 1083 proteins reliably identified either in the cercarial bodies or in microdissected acetabular glands (Supplementary table S15, sheet A). It is obvious from the table that not every gland protein identified in a separate experiment (Supplementary table S11) was detected in the bodies. For the completeness of the results, we also present the re-evaluated MS data of selected samples from the staining experiment, including 548 proteins from cercarial bodies which did not meet our criteria of reliable identification (Supplementary table S15, sheets B and C).

Several proteins which attracted our attention according to their NCBI annotation, and might be involved in invasion or parasite-host interactions, were also identified outside the acetabular glands. These include, e,g., α-taxilin (IL-14), insulin-degrading enzyme, galectin, VAL-6 protein, δ-aminolevulinic acid dehydratase, peptidases and, importantly, a hyaluronidase.

## 4. Discussion

The secretions of cercarial penetration glands that are essential for the invasion of vertebrate hosts have repeatedly been analyzed in various schistosome species. It is obvious that the mode of *in vitro* collection of the gland content and pre-MS sample processing are crucial. Different approaches may result in diverse numbers and spectra of identified proteins (e.g., Knudsen et al., 2005; Mikeš et al., 2005; Curwen et al., 2006; Wang et al., 2006; Dvořák et al. 2008; Hansell et al., 2008). The methods covering mechanically/serum-induced transformation of cercariae, lipid/chemical induction of gland emptying, and skin-induced penetration result in a rather complex mixture of excretory/secretory products from cercariae. These contain mixed secretions from both CAg and PAg, but also other cercarial secretions/excretions, and may contain also significant amount of somatic and tegumental proteins released from transforming, damaged or even dying cercariae, or numerous contaminating proteins from the host skin or complex media. Although some authors observed unparallel release of the contents from particular types of schistosome penetration glands (Haas et al., 1997), according to our experience both types of glands are emptied simultaneously *in vitro* (Mikeš et al., 2005; Řimnáčová et al. 2017). This condition hinders a separate analysis of CAg and PAg secretions. Therefore, we employed the technique of laser microdissection that largely circumvented the above-mentioned drawbacks of the methods used so far for the collection of the contents of penetration glands, and enabled a robust differential proteomic analysis.

For detailed proteomic studies of specific tissues, it is often necessary to deal with a major obstacle - tissue heterogeneity. However, despite the accuracy and effectiveness of laser-assisted microdissection, the proteomic analysis of specific tissues still faces certain limitations, such as the inability to amplify proteins before analysis. Therefore, a precise knowledge of the minimal/optimal amount of material is very advantageous, especially when the target material is available in limited amounts. This was also our case, since the penetration apparatus in a schistosome cercaria consists of only 4 and 6 gland cells, namely CAg and PAg, respectively. Therefore, we decided to test (by using whole cercarial bodies excluding tails first) the effects of the sample preservation methods, amount of material, mode of fixation and staining, and time of processing on the results of proteomic analysis. Based on the cost-per-performance ratio (processing speed/number of identified proteins), we have chosen the volume of 7.5 × 10^6^ μm^3^ of microdissected material per sample and the IST method to process the microdissected penetration glands prior to MS in subsequent principal experiments.

Another factor that has been evaluated was the effect of fixation of cercariae on MS results. Standard histological processing of tissues may potentially give rise to artifacts. According to some authors it is best to use frozen (unfixed) tissue sections for proteomic analysis (Murray, 2007). It is also advisable to incorporate the samples in a cryoprotectant before freezing, such as the optimal cutting temperature medium (OCT) (Bevilacqua and Ducos, 2018). In some cases, however, it is necessary to fix the samples first, e.g., tissues that degrade rapidly. Here we compared non-fixed (NF), ethanol-fixed (EtOH), and (para)formaldehyde-fixed samples (PFA), all frozen after adding the cryoprotectant. Although the use of formaldehyde during tissue fixation leads to cross-links between proteins and other biomolecules (e.g. Ahram et al., 2003; Magdeldin and Yamamoto, 2012), it has been shown several times that similar proteomic data can be obtained from formalin-fixed + paraffin-embedded and snap-frozen samples (Sprung et al., 2009). Although the NF samples provided slightly better results in our study in terms of both the number and the abundance of some proteins when compared to EtOH and PFA-treated ones, all three types of tissue preservation were comparable as for the vast majority of reliably identified proteins. The main advantage of using non-fixed samples was the minimization of product leakage from penetration glands due to tissue shrinkage after adding the fixatives and the best clarity of gland positions in sectioned cercarial bodies (see Fig. 2). Therefore, the use of non-fixed samples seemed to be the best solution in our case.

A precise dissection of very small objects, such as the cercarial penetration gland cells, takes much time. Moreover, the microdissection is performed at RT. Therefore we also tested the effect of the time of exposure to RT on MS results using microdissected cercarial bodies first. We found that even 24h exposure had no considerable effect on any of the captured protein categories. This may be owing to the dessication of cryosections while being stored at –80 °C for ca. 2 weeks prior to microdissection. The stability of samples at RT was another reason why we decided to perform microdissection of gland cells using non-fixed samples.

Finally, we needed to find a way how to distinguish between the two gland types in cryosectioned cercariae. Both of them evince a marked granularity and light refraction that enables to recognize them among other cells on unstained cryosections under a light microscope. Although particular gland types possess distinctive granularity which is well distinguishable by transmission electron microscopy, this was not a reliable marker for light microscopy on cryosections. Therefore, we tested the application of histological dyes. However, the use of some dyes in laser microdissection has been shown to have negative impact on subsequent proteomic analysis. This is known for several commonly used stains, e.g., cresyl violet, eosin, and toluidine blue (Banks et al., 1999; Craven and Banks, 2001; Craven et al., 2002; Moulédous et al., 2002; Ahram et al., 2003). We could not employ navigated laser microdissection (Moulédous et al., 2003) because of the small size of cercariae and associated high risk of cross-contamination of CAg and PAg samples when using this approach. To avoid fixation for the above-mentioned reasons, we decided to test histological dyes in conjunction with non-fixed cryosections. Several histological dyes have been known to distinguish schistosome penetration glands, but most of them can only be used after fixation (Stirewalt and Kruidenier, 1961; Bruckner, 1974; Mikeš et al., 2005; Ligasová et al., 2011). Alizarin is an exception, since it is water-soluble and stains CAg intravitally. The use of PAg-specific ethanol-soluble lithium carmine on non-fixed cryosections of cercariae was not possible since parts of parasite bodies were washed away from the slides. Application of lithium carmine on ethanol-fixed cryosections gave poor staining results. Staining of PAg on cryosections by adding a diluted solution of lithium carmine 1:4 with 70% EtOH gave promissing results in terms of distinguishing the cells. However, this type of staining affected negatively the results of following proteomic analysis. Trypan blue stained cercarial tissues with no specificity, therefore it was not useful for isolation of the glands. Besides, the results of subsequent proteomic analysis showed relatively large number of differentially abundant proteins compared to unstained sections. Also toluidine blue negatively affected proteomic analysis, which is concordant with previous observations of Moulédous et al. (2002). On the other hand, alizarin appeared to be compatible with MS analysis. Thus, we were able to distinguish alizarin-stained CAg from non-stained PA gland cells on non-fixed cryosections, which enabled reliable microdissection. The total proteome of *T. szidati* cercarial bodies (heads excluding tails) resulting largely from the optimization experiments supplements the starring proteomic data obtained from penetration glands. Since it was artificially compiled from two types of separate experiments, it is rather impracticable to interpret the data at a quantitative level. The set of presented proteins includes 1631 identifications in total, but it would be a little bit higher, provided we have included additional non-redundant proteins identified in other optimization experiments (however, they are still present in supplementary tables accompanying these experiments). Our finding is close to the number of MS-identified proteins in whole cercariae of *S. japonicum*, i.e., 1894 including proteins of tails (Liu et al. 2015). The fact that not every protein identified during differential analysis of *T. szidati* glands was detected in whole cercarial bodies can likely be explained by a relatively higher proportion of specific gland proteins in the gland samples compared to the samples of the bodies, because the volumes of both types of samples were the same, and least abundant gland proteins may have not been detected in a more complex samples of total body proteins. The number of assembled cercarial transcripts resulting from a previous transcriptomic analysis of *T. szidati* (Leontovyč et al. 2019) was 8905; that means that around 20 % of proteins potentially present in cercariae were captured in the current study. Moreover, we indentified more than 160 proteins for which the corresponding transcripts did not occur in mixed transcriptomic data of *T. szidati* cercariae/schistosomula (Leontovyč et al. 2019), and their annotations were based on hits in *T. regenti, S. mansoni* or *S. japonicum* genomes. The samples of cercarial bodies were also microdissected from the cryo-sectioned parasites in optimization protocols and attention was paid not to dissect surrounding empty areas of embedding medium, i.e., the dissecting laser was navigated tightly to the surface of microdissected objects. Therefore, it is fair to note that, perhaps, the samples might have partially been depleted of glycocalyx or tegumental proteins.

Utilizing the methods optimized with whole cercarial bodies, we proceeded to a differential proteomic analysis of the contents of *T. szidati* penetration glands. The total number of reliably identified proteins which were found in at least two samples of the triplicate from each gland type was 461. Moreover, other 331 proteins designated as non-reliably identified were detected in one of the replicates. This categorization was performed because of an assumption that in the case of the latter group there may be a higher risk of contamination from surrounding cercarial tissues. Compared to other studies dealing with proteomic analysis of induced secretions of schistosome cercariae, we obtained the highest count of putative penetration gland proteins so far. This could be related to the amount of starting material used, which is in the other studies specified rather vaguely (Curwen et al., 2006; Hansell et al., 2008) or even not specified (Knudsen et al., 2005; Dvořák et al., 2008), and therefore incomparable. There are also questions whether the glands are fully emptied during artificial transformation of cercariae, especially by vortexing or passage through an injection needle, and whether both gland types are emptied equally or not. Moreover, it has been observed that cercariae of *S. japonicum* release lower amounts of secretory vesicles than *S. mansoni* when being induced in serum-free schistosome culture medium (Dvořák et al., 2008). Although the glands are holocrine (Dorsey, 1975) and therefore their whole content including secretory vesicles and cytoplasm should be released, we do not know exactly if this happens under *in vitro* conditions and if the gland content is homogenous in terms of the distribution of vesicles likely carrying a varied cargo. Probably the way of collection of gland secretions most nearing the *in vivo* process was that applied by Hansell et al. (2008) who used skin grafts to induce penetration behavior of *S. mansoni* cercariae. Nevertheless, it is obvious from the study that the time of collection of the products is reflected in the obtained spectrum of peptides/proteins. In the current study, however, we have worked with microdissected gland cells and well defined sample volumes of 7.5 mil. µm^3^ in triplicates. So, it is likely that we had a chance to catch also the low abundant proteins in our shotgun proteomic analyses. It is necessary to mention that in-gel digestion was employed in the older studies and sometimes just visible Coomassie-or silver-stained protein bands or spots after electrophoresis were subjected to MS (Knudsen et al., 2005; Curwen et al., 2006), thus providing data on the most abundant proteins only.

We expected that CA and PA glands of *T. szidati* differ markedly in their protein composition. There was almost equal number of reliably identified proteins in particular cell types (421 in CAg vs. 455 in PAg). Among them, only a minority was exclusive for a particular gland type (6 for CAg and 40 for PAg). Thinking about possible cross-contamination during the microdissection, we suppose that there is a higher risk of contamination of CAg samples with proteins originating from PAg than the other way round. This relates to the arrangement of cytoplasmic projections of the gland cells forming gland ducts. While the cell bodies of PAg are located posteriorly, relatively isolated from CAg, their wide, massive projections line intimately the dorsal surface of CA cell bodies (Ligasová et al., 2011) and might have been unrecognized during microdissection of stained CAg in certain position of cercaria towards the observer. This may be supported by the finding that PAg are more rich in exclusive proteins than CAg. A similar situation could be seen in the case of proteins which were significantly more abundant in one of the two gland types (12 in CAg vs. 48 in PAg). Besides the generally higher prevalence of exclusive and significantly more abundant proteins in PAg compared to CAg, the subsequent KEGG functional annotation revealed that in the PAg there is also more extensive enzymatic equipment in terms of exclusive and significantly more abundant enzymes (29 % vs. 5 %). This holds true especially for hydrolases. Among them, the highest difference in favor of PAg could be observed in case of peptidases and acid anhydride hydrolases (such as GTPases). Thus, this implies that PAg are more catalytically active than CAg in terms of peptido-/proteolysis, and may be indicative of extensive protein synthesis/translocation or vesicle formation/transport in PAg. This was further supported by Gene Ontology enrichment analysis. On the other hand, cysteine-rich secretory proteins (CRISPs) annotated as glioma pathogenesis-related proteins (GLIPRs) or venom allergen-like proteins (VALs) were conspicuously abundant in CAg.

Schistosome penetration gland peptidases have been in focus in the context of the invasion of host skin (histolysis) and other parasite-host interactions. In PAg of *T. szidati*, the cysteine endopeptidase cathepsin L (SmCL3-like) was the fifth most abundant exclusive protein according to intensity score. Its orthologs are expressed in various stages throughout the life cycle of *S. mansoni* and *S. japonicum*, although the highest levels of mRNA detected by qPCR occurred in adult worms in both species and in liver eggs of the latter (confirmed also at protein level) (Dvořák et al., 2009; Huang et al., 2020). Both SmCL3 and SjCL3 were immunolocalized in gastrodermis of schistosomula and adult worms, and in eggshell and adult worm tegument of *S. japonicum*, indicating involvement in blood digestion and action at the parasite-host interface. The localization and abundance of CL3 in PAg of *T. szidati* strongly suggests its involvement in host invasion by schistosome cercariae. The enzyme was probably detected in *S. mansoni* PAg (Dalton et al., 1997) by use of most likely cross-reacting antibodies raised against SmCL2. Cysteine cathepsin F was one of the least abundant exclusive endopeptidases in *T. szidati* PAg. Its orthologue in *S. mansoni*, originally described as SmCL1 (Dalton et al., 1996; Park et al., 2001) was confirmed by RT-PCR and immunoblotting in cercariae (Brady et al., 2000) and is also associated with the gut of adult schistosomes and other helminths (Caffrey et al., 2018). It has also been found in protein extracts of *S. japonicum* cercariae (Liu et al., 2015), but neither CL3 nor CF have been identified by MS in cercarial secretions of *S. mansoni* and *S. japonicum* (Knudsen et al., 2005; Curwen et al., 2006; Dvořák et al., 2008; Hansell et al., 2008). The only exclusive peptidase in *T. szidati* CAg was the cercarial elastase (CE), a clan PA family S1 serine endopeptidase. This enigmatic enzyme has been described from several schistosome species and used to be considered the principal cercarial penetration peptidase and the only serine peptidase involved in degradation of dermal elastin (Salter et al., 2000; Dvořák and Horn, 2018). Several orthologs occur in *S. mansoni, S. haematobium*, and *Schistosomatium douthitti*, suggesting gene duplication after speciation of *Schistosoma*. Two of the eight isoforms in *S. mansoni*, SmCE1a and SmCE1b, were encoded by the most abundant CE transcripts and accounted for over 90 % of the activity (Salter et al., 2002; Ingram et al., 2012). Inconsistent results and discussions have come out concerning the localization of the elastase(s) in a specific gland type (Marikovsky et al., 1990; Bahgat et al., 2001; Salter et al., 2002), and the presence of elastase in *S. japonicum* and in avian schistosomes of the genus *Trichobilharzia*, where no such enzyme has reliably been confirmed until sequencing their genomes (Bahgat and Ruppel, 2002; Mikeš et al., 2005; Dolečková et al., 2007; Dvořák et al., 2007; Kašný et al., 2007; The *Schistosoma japonicum* Genome Sequencing and Functional Analysis Consortium, 2009; International Helminth Genomes Consortium, 2019; https://parasite.wormbase.org). Interestingly, *S. japonicum, T. regenti*, and *T. szidati* posses only one gene coding for an ortholog of the elastase isoform 2b (*S. mansoni* terminology), which comprised the smallest portion of CE transcript in *S. mansoni* sporocysts containing developing cercariae, and was not detected in the form of a protein in *S. mansoni* cercariae (Ingram et al., 2012). In *S. japonicum*, it was a part of cercarial proteome and was immunolocalized in (unspecified) acetabular glands of cercariae (Liu et al., 2015). Although its transcripts occurred in life stages throughout the life cycle, the protein product was only found in sporocysts and cercariae (Huang et al., 2007). Here we proved that in *T. szidati*, the localization of the enzyme was convincingly circumacetabular, which is consistent with the localization of SmCE1a (Dvořák et al., 2008). Its abundance in *T. szidati* CAg was comparable to that of cathepsin L3 in PAg. The previous failure to detect an elastase in *Trichobilharzia* was probably caused by the use of antibodies or degenerate primers based on SmCE1a that is structurally distant from the CE2b ortholog (Mikeš et al., 2005; Dolečková et al., 2007; Dvořák et al., 2008).

Cathepsin B2 (CB2) has been discussed in context of diverse proteolytic tools employed for penetration by cercariae of particular schistosome species (e.g., Dvořák et al., 2008; Dolečková et al., 2009). It was repeatedly localized in PAg of *T. regenti* by immunohistochemistry and immunogold labeling (Dolečková et al., 2009; Ligasová et al., 2011), but in the current study it was identified in both gland types (probably more abundant in PA, but under the level of statistical significance). In *S. mansoni*, it is abundant in tegumental structures of adult worms (Caffrey et al., 2002). Its activity was present in both *S. mansoni* and *S. japonicum* cercarial gland secretions, being ca. 40x higher in the latter (Dvořák et al., 2008). SjCB2 was immunolocalized in cercarial acetabular glands without closer specification of a gland type. However, there were two areas of different immunofluorescence intensity in *S*. j*aponicum* cercariae, probably corresponding to PAg (higher intensity) and CAg (lower intensity) (Zhu et al., 2020). This would correspond with our result. We also showed that TsCB2 was more abundant in *T. szidati* glands than the elastase. The above-mentioned facts stress its importance in the host invasion by *S. japonicum* and *Trichobilharzia* compared to *S. mansoni*, where elastase orthologs seem to be the major histolytic enzymes. Besides the facilitation of skin invasion, TrCB2 and SjCB2 may interfere with functioning of host’s immune response by degrading immunoglobulins and some other immunity-related proteins (Dolečková et al., 2009; Zhu et al., 2020).

The overall most abundant peptidases that occurred almost equally in both PAg and CAg of *T. szidati* belong to metallo(endo)peptidase catalytic type. The leishmanolysin-related invadolysin (family M08) and nardilysin (family M16) were described as essential enzymes in both vertebrate and invertebrate cells, being also secreted by certain cell types and present in body fluids. They seem to be moonlighting peptidases involved in many intracellular and extracellular processes (Di Cara et al., 2013; Kimura et al., 2017; Morita et al., 2017; Abhinav et al., 2019). In *Leishmania* protists, the leishmanolysins have been considered important virulence factors involved, *inter alia*, in degradation of extracellular matrix (e.g, Kulkarni et al., 2008). In *S. mansoni*, leishmanolysins constitute an expanded peptidase family (Silva et al., 2011) and were the second most abundant protease type after elastase isoforms in *S. mansoni* mixed cercarial gland secretions (Curwen et al., 2006). They are also abundant in *S. japonicum* cercarial proteome (Liu et al., 2015) but, interestingly, were not detected in *S. japonicum* cercarial secretions (Dvořák et al., 2008). Although the function of schistosome cercarial leishmanolysins have not been elucidated yet, they are generally believed to be important histolytic tools. In *T. szidati*, the abundance of invadolysin highly exceeded the values obtained for the elastase. Moreover, other three leishmanolysin-like peptidases occurred among non-reliably identified proteins (found just in one replicate in either gland type). This emphasizes the richness and importance of M08 family peptidases in *Trichobilharzia* penetration glands and indicates that they are possibly the most potent invasion tool of cercariae. The M16 family schistosome nardilysin is still a neglected and enigmatic peptidase, probably detected in *S. japonicum* cercarial secretions (Dvořák et al., 2008). Considering it is the second most abundant peptidase in *T. szidati* glands, it might broaden the spectrum of potential histolytic metallopeptidases.

Conspicuous was the wide representation of dipeptidases (DPP) and oligopeptidases (OPP) which were either significantly more abundant in PAg or present in both gland types of *T. szidati* without significant difference. They are mostly localized intracellularly and have been considered to play general housekeeping roles in cells of virtually all tissues in eukaryotes. For example, the serine DPP2 is a lysosomal aminopeptidase, although it has been found also in cytoplasm, vesicles and body fluids (Maes et al., 2007; Prajapati and Chauhan, 2011). The metallo DPP3 is generally a cytosolic aminopeptidase, also secreted into the lumen of some organs (Chen and Barrett, 2004; Prajapati and Chauhan, 2011). Their roles in various cell types have not been fully elucidated. Based on their ability to catabolize oligopeptides, their involvement in terminal stages of protein turnover, activation of other proteins, and inactivation of bioactive peptides such as angiotensin, substance P, bradykinin, and some hormones have been confirmed or suggested. This points to their implication in blood pressure regulation, inflammation and pain modulation which may be useful for the invading parasite. We identified three DPP3 and two DPP2 in the glands of *T. szidati*. Although these peptidases occurred also in cercarial proteome of *S. japonicum* (Liu et al., 2015), it looks strange in the view of our results that they have not been identified in induced gland secretions of *S. japonicum* and *S. mansoni* (Knudsen et al., 2005; Curwen et al., 2006; Dvořák et al., 2008; Hansell et al., 2008). Thus, their involvement in the invasion of host skin by cercariae and parasite-host interactions *in vivo* remains unclear. The same can actually be applied to the prolyl and thimet oligopeptidases, leucyl aminopeptidase, and the PAg-exclusive serine carboxypeptidase cathepsin A. These are ubiquitous in cells of eukaryotes where they make up a constitutive peptidolytic equipment participating in, e.g., degradation of peptides generated by proteasomes, peptide catabolism in digestive cells, and are frequently mentioned in context with the catabolism of bioactive peptides such as bradykinin, angiotensin, substance P, and endothelin 1 (e.g., Saric et al., 2004; Timur et al., 2016; Fülöp et al., 2020). This implies the interference of schistosome cercariae with blood pressure and vasoconstriction at the site of parasite entry to the circulatory system. Although some of these enzymes have also been found or studied in various life stages of schistosome parasites (e.g., McCarthy et al., 2004; Fajtová et al., 2015; Liu et al., 2015), we bring the first direct evidence of their presence and relatively high abundance in cercarial penetration glands. Also, the interesting first finding of leukotriene A4 hydrolase in schistosome penetration glands brings another view on how penetrating schistosome cercariae may be able to effectively affect the function of host’s immunity. This enzyme is a bifunctional protein possessing both epoxide hydrolase activity converting leukotriene (LT) A4 to LT B4 and metallo aminopeptidase activity that is just believed to be involved in processing of bioactive peptides related to inflammation and immune defense (Haeggström, 2004). It was more abundant in both gland types than, e.g., particular cysteine cathepsins and elastase. However, its role in schistosome-host interactions remains to be elucidated, since LT B4 possesses pro-inflammatory and T-cell-attracting properties, that, at first sight, do not seem to be advantageous for the parasite.

The high abundance of two peptidase inhibitors of two classes, a serpin superfamily alpha-1-antiproteinase-like protein and cystatin B (cytoplasmic stefin), implies that the proteolytic environment in the glands may be strictly regulated. Interestingly, while the equal presence of cystatin B in both gland types corresponds with the localization of TsCB2, the serpin is significantly more abundant (39 fold change) in PAg than in CAg, where the elastase has exclusively been localized. Similarly, endosomal cystatin F, a cathepsin C-directed peptidase inhibitor (Hamilton et al., 2008), was found in both gland types, while its target was detected exclusively in CAg (but only in one of three replicates, which might have distorted the interpretation). We are not aware of a study reporting on the presence of cystatins in cercarial gland secretions except for *S. japonicum* (Dvořák et al., 2008). Serpins have been detected in gland products of *S. mansoni* cercariae (Curwen et al., 2006; Hansell et al., 2008). Thus, when being secreted from the glands, we cannot exclude the involvement of these inhibitors in defense against the action of host’s peptidases utilized by immune cells, or as anticoagulants or complement suppressors in the case of serpins (e.g., Sugino et al., 2003; Hamilton et al., 2008; Molehin et al., 2012; Nanut et al., 2017).

Since the synthesis of various peptidases and peptidase inhibitors localized in *T. szidati* penetration glands can be expected also in other cells of cercariae (e.g., general house-keeping cytoplasmic and lysosomal proteins), and the transcription of coding genes may be specifically regulated in different cells, the comparison of the abundances at the protein and mRNA levels may be somewhat misleading. Here we presented proteomic data on isolated penetration glands, while *T. szidati* transcriptome was reconstructed using whole cercarial bodies including tails (Leontovyč et al., 2019). Nevertheless, in some cases we can discuss the recurrent question concerning proteosynthesis in free-swimming matured cercariae and in embryonal cercariae developing from germ balls inside the intramolluscan stages, the sporocysts. For instance, as can be seen in Fig. 13, for three abundant gland proteins (alpha-1-antiproteinase, invadolysin, and elastase), there were virtually no transcripts detectable in cercarial transcriptome. This strongly implies that these proteins are synthesized in pre-emergent developing cercariae to be ready in the glands at the time of contact with the host. On the other hand, the relatively high abundance of some transcripts if compared to the low abundances of the proteins within the glands (e.g., calpain, cystatins, cathepsins A and F) may suggest that their genes are transcribed in many other cercarial cells, perhaps even at a higher level, or that their transcription (or translation of mRNA pre-synthesized in intramolluscan stages) occurs also in the glands of free-swimming cercariae. In this case, we cannot provide an acceptable answer, although transcriptomic analysis of microdissected gland cells would enable due considerations.

Besides peptidases and their inhibitors, schistosome SCP/TAPS domain-containing proteins, including VAL or GLIPR-like proteins (depending on annotation), seem to be important for interactions of the penetrating cercariae with the host (e.g., Kelleher et al. 2014; Fernandes et al. 2018). In *S. mansoni*, the highly expanded family of VAL proteins includes 28 members, of which 23 possess a signal peptide predestining them for secretion (Chalmers et al. 2008; Philippsen et al. 2015). VAL4, VAL10, VAL18, and VAL19 were detected in induced cercarial secretions or in CAg of this species (Curwen et al. 2006; Hansell et al. 2008; Farias et al. 2019). In *T. szidati*, the picture seems to be similar. An ortholog of SmVAL18 (Tszi_020665|m.170901 annotated as GLIPR-1) was the most abundant exclusive protein in CAg. Moreover, other SmVAL orthologs appeared significantly more in CAg of *T. szidati*: VAL4 (Tszi_003791|m.45920), VAL18 (Tszi_023061|m.180336 also annotated as SjVAL5), and VAL10 (Tszi_020765|m.171327 and TRE_0000208601-mRNA-1 annotated as GLIPR-1). We did not detect a VAL19 ortholog in *T. szidati* glands. However, the terminology of VALs seems to be somewhat confusing among different schistosome species and we used always the first relevant term from NCBI annotation in the tables. Therefore, a more precise evaluation of the set (including highly similar isoforms?) of *T. szidati* VALs detected in this study would require a deeper analysis based on combination of (unavailable) genomic data, transcriptomic data, and alignments/phylogenetic analysis in context of VALs from other schistosome species, which is beyond the intention of this paper.

Interesting was the finding of a DM9 domain-containing protein that was highly abundant in PAg. This domain has been found in a variety of both vertebrate and invertebrate proteins, sometimes fused with other domains possessing cytolytic activities, such as in natterin-like toxins (Lopes-Ferreira et al. 2014; Jia et al. 2016). They have also been characterized as pattern-recognition receptors with agglutinating activities and a range of recognition spectrum towards lipopolysaccharide, peptidylglycan, mannan, and β-1,3-glucan. Their involvement in innate immunity is highly expected (Jiang et al. 2017). These features of DM9 domain-containing proteins could possibly attribute to a protein with lectin activity previously localized in *T. szidati* PAg (Horák et al. 1997), although the sequence of this lectin is not known. We can only speculate on a possible involvement of the DM9-domain protein in interactions of penetrating cercariae with host’s immunity or in cytolysis. Similarly, the LAMA-like protein 2 ortholog that was highly abundant and exclusive in PAg might contribute to cercarial invasion of host’s skin and underlying tissues due to phospholipase-B activity, which may aid to damage of host cell membranes as has been proven in, e.g., pathogenic yeast (Leidich et al. 1998). It is virtually pointless to speculate on biological functions of many of the proteins found in the glands. They are either largely unknown or would be just inferred from their mammalian homologs (provided that functional studies are available). This may lead to false interpretations, since vast differences in structure may result in varied biological activities. Instead, we would like to point to some important and yet not reliably answered questions concerning the protein composition of schistosome penetration glands and the process of cercarial penetration. For example, the holocrine nature of the glands predetermines the whole content, including vesicles and cytoplasm, to be secreted. However, there is a discrepancy between the results of our proteomic analysis of microdissected glands and the results of similar analyses performed with isolated gland secretions of other schistosome species. The secretions induced by different methods (see above) have been always conspicuously poorer in the spectrum of identified proteins compared to gland proteomes in the current study. Even some highly abundant proteins recognized in *T. szidati* glands have not been detected in, e.g., *S. mansoni* and *S. japonicum* secretions. A significant number of them, including some acid peptidases and glycosidases, are known from lysosomes of other animal cells. To our best knowledge, there have been no studies so far performed on the qualitative composition of schistosome gland vesicles - are they varied within a particular gland type? Also, the presence of “true” lysosomes within the glands and their exocytosis have not been studied (Tancini et al. 2020). Thus, besides the few examples of gland peptidases studied in the context of histolysis of host tissues, such as elastases, cathepsin B2 and leishmanolysins, uncovering the functions of many others requires further research. Moreover, it is not clear if the (potentially varied) secretory vesicles are evenly distributed within the glands. In this work, we were only able to dissect gland cell bodies; their cytoplasmic projections (“ducts”) stuffed with vesicles had to be omitted during microdissection, since the duct bundles are formed by parallel CAg and PAg lines (Ligasová et al. 2011) and cross-contamination would certainly occur. This fact might have possibly affected our results towards the real situation. Similar questions apply also to many cytosolic proteins from the glands.

Furthermore, there are some other unanswered questions concerning the compartmentalization of schistosome penetration apparatus into two gland types rapidly differing in their composition. In *in vitro* studies, the contents of both PAg and CAg were emptied simultaneously upon the stimulation by a general signal, the unsaturated fatty acids such as linoleic and linolenic acids (e.g., Mikeš et al. 2005; Řimnáčová et al. 2017). On the other hand, lipid fractions and hydrophilic extracts from the skin stimulated gland emptying (partly) differentially *in vitro* (Haas et al. 1997). However, we still cannot be sure what is the situation *in vivo*. Therefore, there is also a possibility that the two compartments arose in the course of evolution from the need to separate certain compounds which are then cross-activated while being mixed upon release of gland contents. Unfortunately, we still do not know anything about the conditions necessary for activation of gland peptidases which are produced in the form of pro-enzymes, e.g., the involvement of some of them in the trans-processing of the others. Also, it is obvious that pH values at which specific groups of the gland peptidases express their (optimal) activities greatly differ. So, what are the mechanisms ensuring optimal pH for the action of particular peptidases during cercarial penetration throug the layers of the skin? And, finally, is this compartmentalization at the level of gland cell types typical of schistosomes as a consequence of their adaptation to the warm-blooded vertebrate hosts? Regrettably few in-depth studies were in fact performed regarding the ultrastructure and biochemical composition of penetration glands of other trematode cercariae; for example, in the case of a fish skin-invading species *Diplostomum pseudospathaceum* the four cercarial penetration gland cells appear morphologically uniform (Moczoń 1994).

In conclusion, we present here a thorough and the most detailed insight into the protein contents of schistosome cercarial penetration glands so far, employing up-to-date techniques of laser microdissection and tandem mass spectrometry. We believe that the differential proteomic analyses of the two gland types greatly enhanced the knowledge of general composition of the glands and quantitative representation of the proteins, and moved forward the knowledge of localization of individual protein species within particular gland types. Although the contamination of the specific samples by proteins originating from the other gland type or from surrounding tissues cannot be completely excluded, we attempted to optimize procedures of sample collection which should minimize such a possibility. Besides, we present for the first time complex proteomic data obtained from an avian schistosome. Our results also induced some additional yet unresolved questions which established perspectives for further investigation of proteins and processes involved in skin penetration and early parasite-vertebrate host interactions by schistosome cercariae.

## Acknowledgements

The authors would like to thank Professor Pavel Dundr and Dr. Ivana Stružinská (Institute of Pathology of the First Faculty of Medicine, Charles University and General University Hospital, Prague, Czechia) for the possibility to use the device for laser microdissection, and Veronika Siegelová (Faculty of Science, Charles University, Prague, Czechia) for great maintenance of the parasite life-cycle.

## Authors’ contribution

PH, JB, and OV designed the study. PH supervised the study. OV prepared material and performed laser microdissections. PT designed procedures of sample preparation for LC-MS/MS and performed MS analyses. OV, LM, PT, and RL contributed to analyses of the data. RL provided and processed transcriptomic data. OV and JB prepared the pictures. OV and LM interpreted the findings, drafted and finalized the manuscript. All authors contributed to writing and approved the final version of the manuscript.

